# Coordinated inhibition of SOX9 and cell cycle progression by microRNA-200 restricts sebaceous gland fate specification

**DOI:** 10.64898/2026.01.09.698672

**Authors:** Arpan Das, Yuheng C Fu, Haimin Li, Megan A. Wong, Annalina Che, Anumeha Singh, Jimin Han, Glen Bjerke, Dongmei Wang, Rui Yi

## Abstract

The microRNA-200 family (miR-200s) is widely recognized for their potent role in inhibiting epithelial-to-mesenchymal transition and cell cycle progression in cancer. However, their functional specificity in normal epithelial development remains poorly understood. Here we show that miR-200s are highly enriched in hair matrix progenitors but conspicuously absent from the upper hair follicle (HF), the anatomical location where sebaceous gland (SG) and HF stem cells are specified. We demonstrate that elevated miR-200 expression in this region abolishes SG fate specification while permitting hair morphogenesis. Genome-wide identification of miR-200 targets reveals that miR-200s regulate multiple negative regulators of WNT signaling, in addition to cell cycle regulators. Single-cell and spatial transcriptomic analyses uncover a mutually exclusive expression patterns between WNT activity and SOX9 in the upper HF, which is disrupted by miR-200 induction, resulting in compromised SOX9 function. Mechanistically, genome-wide identification of SOX9 targets uncovers a broad network of lipid and fatty acid metabolism genes critical for the transition of upper HF progenitors to the SG fate. The coordinated inhibition of the SOX9-dependent lipogenic program and the potent restriction of cell cycle progression, both mediated by miR-200s, collectively blocks SG specification. Taken together, this work reveals an unexpected specificity of miR-200 in restricting epithelial plasticity and elucidates a spatially defined SOX9 regulatory network essential for SG development.

## Introduction

The miR-200 family of microRNAs (miRNAs) is highly expressed in epithelial tissues and is best known for their potent tumor suppressive functions. In cancer, miR-200s function as potent inhibitors of epithelial-to-mesenchymal transition (EMT) and cell cycle progression^1–4^, therefore restraining metastasis. Mechanistically, gain-of-function studies in metastatic cancer cells and mouse models have established the transcription factors (TFs) ZEB1 and ZEB2, which are core drivers of the mesenchymal fate^5^, as *bona fide* targets of miR-200s. Their downregulation by miR-200s restores epithelial gene expression, such as E-cadherin, and reverts invasive, metastatic phenotypes back toward the epithelial fate^1,4^. However, a paradox remains: miR-200s are predominantly expressed in normal epithelial cells during development and homeostasis, where *Zeb1* and *Zeb2* are largely absent. Instead, experimental capture of miR-200 targets in primary keratinocytes has identified a large number of *bona fide* targets regulating the actin cytoskeleton, cell adhesion, and multiple signaling pathways^6^. Consistent with these findings, functional studies using both miR-200 knockout (KO) and inducible mouse models demonstrate that miR-200s primarily orchestrate actin cytoskeletal dynamics, adhesion, and cell proliferation rather than suppression of classic EMT markers during embryonic skin development^7^. These results raise an important question: how is the intrinsic developmental role of miR-200s in regulating epithelial tissue development related to their potent anti-EMT and tumor suppressor activity in cancer?

To address this question, the morphogenesis of skin appendages offers a powerful paradigm of epithelial plasticity. The transformation of a stratified epithelium into morphologically distinct, self-sustaining mini-organs, such as sebaceous gland (SG) and hair follicle (HF), requires precise spatiotemporal coordination of signaling cues, including the WNT, SHH and BMP pathways. WNT/β-catenin signaling is essential for the initiation of the HF fate and the subsequent differentiation of hair shaft^8,9^, while SHH drives the proliferative expansion of progenitors and hair morphogenesis^9–11^. Conversely, BMP signaling acts to maintain the quiescence of adult HF-SCs and also promotes hair differentiation^12–14^. Intriguingly, the specification of the SG represents a distinct lineage divergence that requires the establishment of a unique niche in the upper HF^15,16^, governed by specific signaling and transcriptional programs. Indeed, genetic perturbations in WNT, BMP and SHH signaling pathways as well as transcriptional programs controlled by various transcription factors (TFs) have been shown to cause defects in SG morphogenesis and development^17–24^. Notably, these signaling pathways and TFs critical for skin appendage formation, such as WNT, SHH, GLI and SOX9 TFs, have been identified as driver pathways for tumorigenesis or maintaining the stemness of tumor initiating cells in squamous cell carcinoma and basal cell carcinoma^25–28^. Thus, the specification and morphogenesis of SG and HF provide a sensitive platform to dissect the impact of miR-200s in regulating cell fate specification.

In this study, we determine the function of miR-200s in regulating epithelial plasticity by studying their roles during skin appendage formation. Surprisingly, induced miR-200 expression in these upper HF progenitors specifically inhibits sebaceous gland (SG) fate specification but permits hair morphogenesis. By combining single-cell RNA sequencing (scRNAseq), spatial transcriptomics and phenotypic analysis, we show that miR-200s inhibit multiple negative regulators of WNT signaling while potently inhibits the cell cycle progression. These actions lead to the inhibition of a SOX9-mediated transcriptional network that activates the SG fate. Together, this work reveals an unexpected specificity of miR-200s in antagonizing SOX9 function and SG lineage commitment during skin appendage formation. These findings provide evidence that the EMT-suppressive activity of miR-200s in cancer likely emerges from their intrinsic role in constraining epithelial plasticity programs such as the SOX9-mediated appendage formation.

## Result

### Expression pattern of miR-200s during skin morphogenesis

The five miR-200 family members (miR-200s), clustered in two genomic loci and sharing similar sequences in both 5’ seed and 3’ complementary regions (Fig. 1a), are highly expressed during embryonic skin development^7^. We previously reported that miR-200b, a representative miR-200 family member, is broadly expressed in basal epidermal progenitors and hair placode during embryonic skin development^7^. To determine the spatial patterns of miR-200s during the formation of skin appendage, such as hair follicle (HF) and sebaceous gland (SG), we performed fluorescence in situ hybridization (FISH) to detect miR-200b together with other epithelial lineage markers in postnatal day 3 (P3) skin when HFs develop into hair peg and more mature HFs. Notably, miR-200b expression was highly enriched in the Matrix progenitors and largely depleted from the upper HF regions and outer root sheath (ORS) (Fig. 1b), where SG progenitors and hair follicle stem cell (HF-SC) progenitors as well as ORS progenitors reside, respectively. These data reveal a spatiotemporally restricted miR-200 expression in skin appendages.

**Figure 1:**
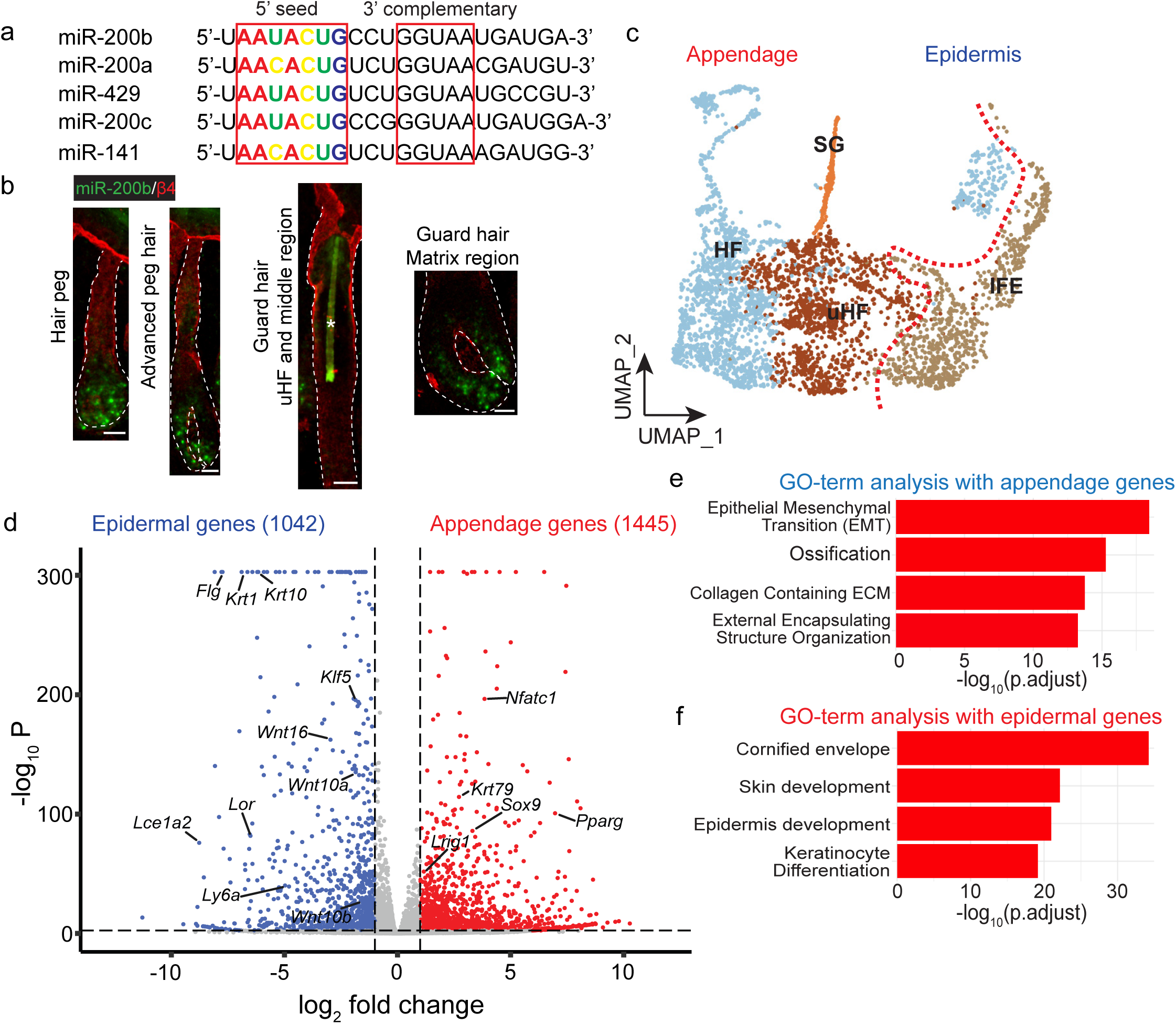
Expression pattern of miR-200s and single-cell analysis of epithelial transcriptome during skin morphogenesis. **a.** All five miRNA sequences from the miR-200 family are shown with the conserved seed sequence highlighted in the left red rectangle. The sequences marked by the red rectangle in right side highlight an additional conserved 3’ sequence. **b**. Fluorescence *in situ* hybridization (FISH) of miR-200b-3p (green) at different HF stages at P3 shows that the expression of miR-200b is restricted to the matrix progenitors. β4-integrin (red) marks the basement membrane. The green autofluorescence signal from the hair shaft is noted by *. Scale bar: 20 µm. **c**. UMAP representation of the epithelial cells of the skin. Four major groups of skin epithelial cells: IFE, uHF, HF, and SG are shown in different colors. Red line marks the boundary between epidermal and appendage-keratinocytes. **d**. Volcano plot of the differentially expressed genes in appendage- and epidermal-compartments, in red and blue, respectively. Selected genes are marked on the volcano plot. **e**. & **f**. GO-term analysis of appendage genes **(e)** and epidermal genes **(f)**, respectively.

Because the miR-200 family is known to inhibit the epithelial-to-mesenchymal transition (EMT), we sought to determine which epithelial cell populations and lineages contain the EMT-like signatures, which could be targeted by miR-200s. To this end, we performed single-cell RNAseq (scRNAseq) at postnatal day 3 (P3) when interfollicular epidermis (IFE), SG and HF were readily distinguishable (Extended Data Fig. 1a). Upon quality control and data processing (see Method), subclustering of epithelial cells identified all cell types of these three lineages (Fig. 1c and Extended Data Fig. 1b-e). We next separated IFE cells from skin appendage cells, including SG and HF cells, and performed differential gene expression (DEG) analysis. As expected, genes associated with IFE, such as *Krt1*, *Krt10*, *Klf5*, and *Flg*, and genes associated with the SG and HF fate, such as *Sox9*, *Nfatc1*, *Krt79*, *Pparg*, were identified as the most differentially expressed (Fig. 1d and Table S1). Many WNT ligands such as *Wnt10a*, *Wnt16* and *Wnt10b*, which are also known to be expressed in hair matrix cells, were more highly enriched in IFE cells, consistent with their production in IFE cells through P63-mediated activation during embryonic development^29^ and their role in maintaining IFE stem cells in adult^30^. Gene ontology (GO) analysis further revealed genes associated with EMT as the most highly enriched in the SG and HF cells whereas epidermal development genes were highly enriched in the IFE (Fig. 1e-f). Notably, *Sox9* was a highly distinct “appendage” gene whereas *Klf5* was a highly distinct epidermal genes, in line with their role as lineage defining master regulators in homeostasis, wound repair and cancer^31^. These results are consistent with the notion that skin appendage formation is an EMT-like process, which undergo extensive signaling and transcriptional changes and cellular remodeling.

### Enhanced miR-200 expression abolishes sebaceous gland formation

To test which skin appendages are regulated by miR-200s, we used an established *Krt14-rtTA/pTRE2-miR-200b/a/429* model to examine the effect of miR-200 expression in skin morphogenesis (Fig. 2a). During key developmental stages of skin morphogenesis from E15 to P3, we observed no strong perturbation to IFE development and hair placode formation (not shown). Strikingly, however, the SG was largely absent in miR-200 induced skin by P3 (Fig. 2b-c and Extended Data Fig. 2a). To determine how the SG formation was inhibited by miR-200, we examined the spatiotemporal induction of PPARγ, the critical transcription factor (TF) for maintaining the SG fate^15,23^. Between E17 and E18, PPARγ first emerged in the suprabasal cells of the upper HF region of guard hair at the hair peg stage. In more mature guard hair in which hair matrix differentiation was initiated, SCD1, a marker for SG maturation^15,32^, was also detected in these suprabasal PPARγ+ cells (Extended Data Fig. 2b). In addition to these suprabasal PPARγ^hi^ cells, a few basal cells in the upper HF region showed weaker PPARγ signals and were SCD1 negative (Extended Data Fig. 2b-c). At these early stages, no PPARγ+ cells were detected in less mature secondary HFs (awl and zigzag) that were still in the hair placode or hair germ stages. At postnatal day 1 (P1), more basal cells of the upper HF region became PPARγ+ when guard HFs further elongated into the dermis, and PPARγ+ cells were universally detected in all HFs including awl and zigzag (Extended Data Fig. 2d). By P3, the SGs were readily observed morphologically and by oil red O staining in all control HFs, regardless of their type (Fig. 2b). In contrast, the SGs were largely absent from all HFs, including both more mature guard hair with well differentiated hair matrix as well as awl and zigzag hair in miR-200 induced mice (Fig. 2d). To distinguish whether miR-200 expression blocks the fate specification of the SG and/or inhibits the expansion and differentiation of SG progenitors after specification, we quantified IF results for PPARγ and SCD1, which mark the progenitors and differentiated SG cells, respectively^15,16^. On average, we observed similarly reduced (∼4-fold reduction) PPARγ+ and SCD1+ cells in induced HFs (Fig. 2e). Because we occasionally observed PPARγ+ and SCD1+ cells in some induced samples, these HFs allowed us to pinpoint how enhanced miR-200 expression inhibits the SG formation. To this end, we examined the correlation between miR-200 expression and PPARγ+ cells by performing miR-200b FISH and PPARγ IF in the same HFs. Notably, in miR-200 induced HFs, PPARγ was absent in all HF cells with highly induced miR-200 expression whereas the few PPARγ+ cells were detected invariably in miR-200 low or negative cells (Fig. 2f). These cells likely reflected mosaic patterns of *Krt14-rtTA-*mediated induction or the selection against miR-200 induced cells, which were less proliferative (see below). In support, when we further examined the expression of QKI, a well-established miR-200 target^7^ and whose expression was strongly repressed in miR-200 induced epithelial cells, PPARγ+ signals were only detected in a few HFs in which QKI expression was not inhibited by miR-200 (Fig. 2g). Thus, both miR-200 expression (FISH) and function (QKI inhibition) were inversely correlated with the appearance of PPARγ+ cells and the formation of SG. Taken together, these genetic and phenotypical results reveal the specificity of miR-200s in potently inhibiting the SG fate specification and SG morphogenesis in a cell-autonomous manner.

**Figure 2:**
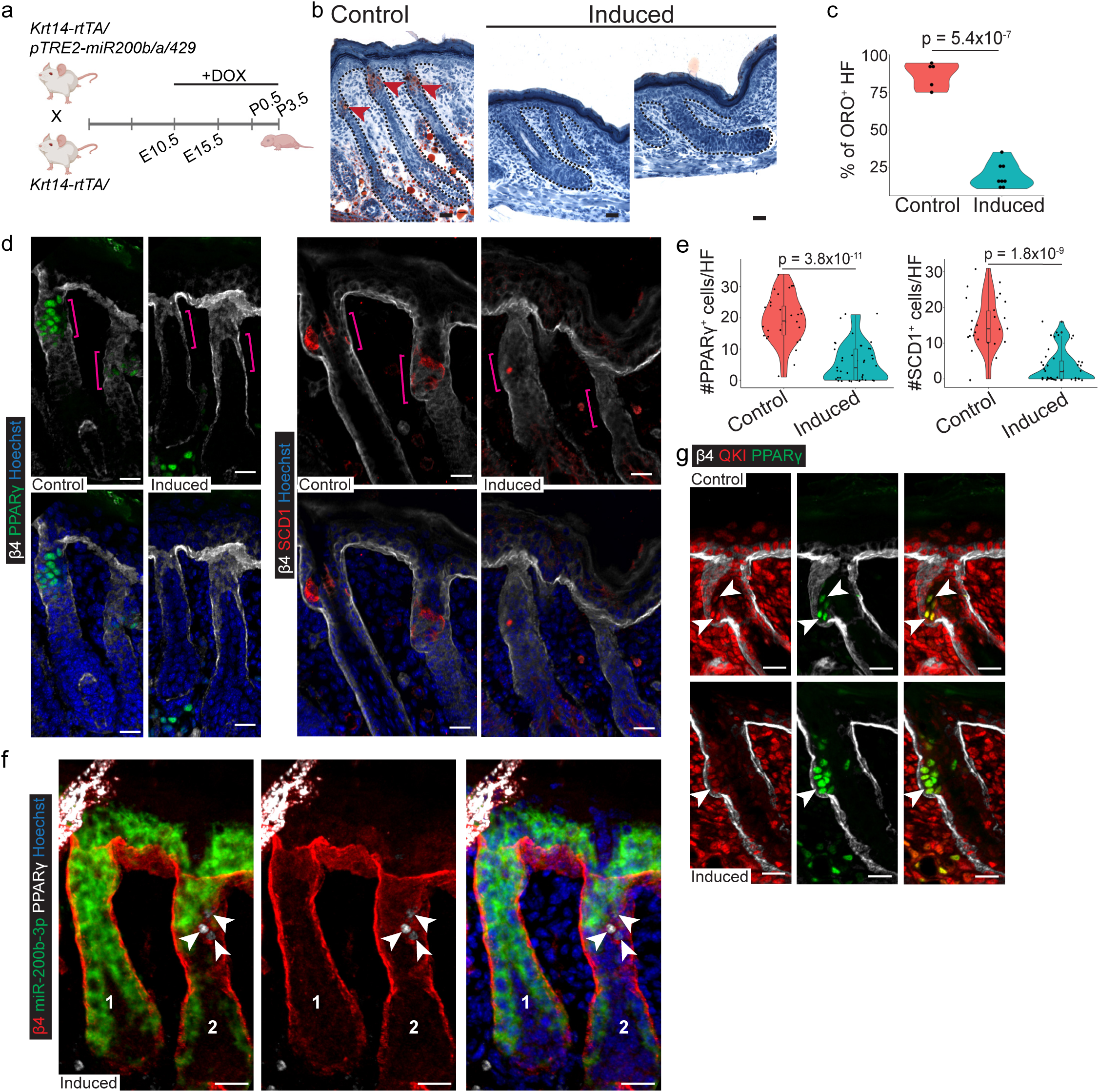
Enhanced miR-200 expression inhibits sebaceous gland specification. **a.** Schematics of miR-200b/a/429 inducible mouse model. **b.** Representative image from Oil Red O staining performed on control and induced dorsal skins at P3. Black dotted lines outline the epidermis and hair follicle, and red arrowheads point to the ORO^+^ SGs. Scale bar: 20 μm. **c.** Quantification of ORO^+^-HFs at the P3 stage between control (N=5 animals, n=66 HFs) and induced (N=8 animals, n=253 HFs) samples. T-test was performed for statistical analysis. **d. & e.** Immunofluorescence (d) and quantification (e) of PPARγ and SCD1 in control (N=3 animals, n=30 HFs) and induced (N=4 animals, n=43 HFs) skin from P3 animals. Magenta brackets in **(d)** indicate the upper HF region. Scale bar: 20 μm. T-test was performed for statistical analysis in **(e)**. **f.** miR-200b FISH and PPARγ IF show cell-autonomous inhibition of sebaceous gland fate by miR-200s. HF #1 with uniform miR-200 induction lacks PPARγ^+^-sebocytes, whereas HF #2 with mosaic miR-200 induction has PPARγ^+^-sebocytes (white arrowheads) that do not have elevated miR-200 expression. Scale bar: 20 μm. **g.** IF for PPARγ and QKI shows the inverse relationship between miR-200 induction and sebocyte specification. Scale bar: 20 μm.

### Single cell trajectory analysis reveals the blocked sebaceous gland specification

To investigate the impact of miR-200 on skin development and appendage formation at single-cell resolution, we performed scRNAseq in miR-200 induced skin and compared the results with P3 control. In control sample, the SG cluster was readily detected by the high expression of *Pparg* and *Scd1* (Fig. 3a-b and Extended Data Fig. 3a-b). Lending further support to the morphological study, only the SG lineage was conspicuously absent whereas IFE and HF cell clusters were largely unchanged in the induced skin (Fig. 3a). We then focused on uHF and SG cell clusters (Fig. 3b) to examine the effect of miR-200 expression on the transcriptome of these progenitors. The epidermal score, calculated by summing the expression of all IFE enriched genes (Fig. 1d), was significantly increased across all of these epithelial cells in induced skin (Fig. 3c), whereas the cell cycle score was significantly reduced across these epithelial cell clusters (Fig. 3c). These data provided orthogonal validation for the strong and specific inhibition of the SG fate specification mediated by miR-200 and identified the repression of cell cycle progression as the global outcome of miR-200 induction in upper HF cells.

**Figure 3:**
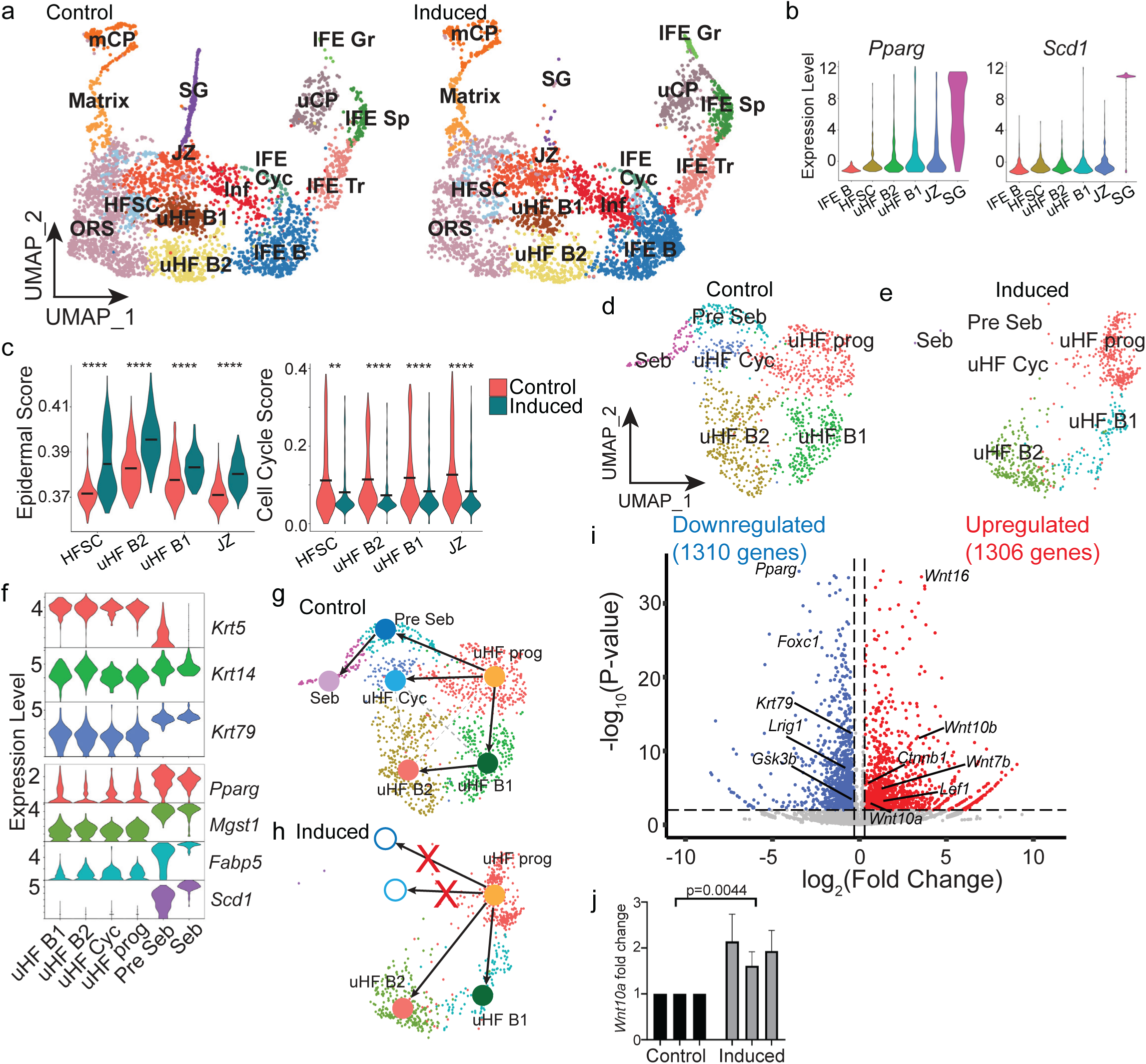
Single cell trajectory analysis reveals the blocked sebaceous gland specification. **a.** UMAP representation of Seurat-integrated epithelial clusters from control (left) and induced (right) dorsal skin at P3. **b.** Violin plots of *Pparg* and *Scd1* expression in epithelial cell clusters from the upper HF regions. **c.** Violin plots comparing the epidermal (left) and cell cycle (right) scores between control and induced HF-SC, uHF B1/B2, and JZ clusters. The Wilcoxon test was performed for statistical comparison. **d & e.** UMAP representations of re-clustered upper HF cells, including uHF B1/B2, JZ, and SG clusters from Fig. 3a, from control (**d**) and induced (**e**) skin. **f.** Violin plots of selected marker genes used to identify the uHF-related clusters. **g & h.** scVelo trajectory analysis of control **(g)** and induced **(h)** uHF and SG cells. The uHF progenitor-to-SG and uHF progenitor-to-cycling cell trajectories were blocked. **i.** Volcano plot of differentially expressed genes between control and induced uHF progenitor cluster. **j.** RT-qPCR analysis of *Wnt10a* in control and induced epidermal cells (n=3 pairs of control and induced samples). T-test was performed for statistical analysis.

We next leveraged these scRNAseq data to define how miR-200 blocks SG fate specification. To this end, we focused on cell clusters corresponding to upper HF regions, including uHF progenitors (uHF prog), two uHF basal-like clusters (uHF B1 and B2), cycling cells in uHF (uHF cyc) and SG (Pre seb and seb) cells (Fig. 3d-f). Among these uHF clusters, key markers for upper HF regions, such as *Lrig1* and *Lgr6*^33–36^ but not *Lgr5*^33,37^, were enriched or depleted as expected, validating our profiling and clustering strategy (Extended Data Fig. 3c). The identification of these clusters in neonatal skin was also in line with a recent single-cell trajectory study of the SG lineage in adult^15^. Comparative trajectory analysis between control and induced samples uncovered a blockage on SG fate specification downstream of uHF progenitors and the complete loss of the cycling cell cluster in uHF (Fig. 3g-h). In contrast to the blocked uHF progenitor-to-SG fate and uHF progenitor-to-cycling cell trajectory, both steady and dynamic trajectory analyses^38^ revealed the intact lineage progression from uHF progenitors to two uHF basal-like populations (Fig. 3g-h), reflecting the specificity of miR-200s for SG fate specification. Because the blocked uHF progenitor to SG transition, we reasoned that gene expression changes in uHF progenitor cells, consisting of progenitors to both uHF basal cells and the SG lineage, should reveal some effect of miR-200 on SG fate specification. Interestingly, DEG analysis of uHF progenitor between control and induced samples revealed that genes associated with canonical WNT signaling, including *Wnt16* and *Wnt10a/b* ligands, *Lef1* TF and *Ctnnb1*, were prominently upregulated in induced samples, whereas *Pparg, Foxc1*, *Krt79*, and *Lrig1*, which were implicated in SG morphogenesis^15,39^, were significantly downregulated in these progenitors (Fig. 3i). To validate these findings, we performed LEF1 and FOXC1 IF staining to confirm the elevated or reduced expression of these key genes, respectively, in E18 embryonic skin when the initial SG specification takes place (Extended Data Fig. 3d-e). Furthermore, quantitative PCR analyses of *Wnt10a* mRNA validated increased *Wnt10a* expression (Fig. 3j). Taken together, these genomic data highlight the specificity of miR-200s in blocking SG fate and reveal dysregulated WNT-related genes in uHF progenitors.

### Perturbed BMP signaling does not abolish sebaceous gland formation

We next sought to determine the mechanism underlying the abolished SG specification caused by miR-200s. Genome-wide capture of miR-200-bound targets by CLEAR-CLIP identified numerous *bona fide* miR-200 targets in the pathway of cytoskeleton and cell cycle regulators^6,7^. Closer inspection of miR-200 interacting mRNAs^6^ and predicted targets^40^, however, revealed several key regulators of BMP, including *Nog*, *Bmpr1a* and *Foxc1*, and negative regulators of WNT, *Csnk1a1* and *Btrc (*see Fig. 5), as miR-200 targets. Because *Nog* overexpression in the skin led to ectopic SG formation along the outer root sheath (ORS) layer and SG hyperplasia^18,19^, these results suggest a potential role of *Nog* in regulating SG specification. We first validated *Nog* as a *bona fide* target of miR-200 through a single, conserved miR-200b/c/429 binding site, matching No. 1-8nt of miR-200, at the 3’UTR (Fig. 4a). Next, we generated a skin specific *Nog* conditional knockout (cKO) model (*Krt14-Cre/Nog^fl/fl^*) to delete *Nog* from epithelial cells of the skin. Similar to miR-200 induced mice, *Nog* cKO skin developed largely normal epidermis and HF (Fig. 4b). Characterization of the SG by oil red O (Fig. 4b), PPARγ+ and SCD1 staining (Fig. 4c) revealed compromised SG formation with the reduced SG size. However, while both oil red O production and SCD1+ mature sebocytes were reduced, PPARγ+ cells were clearly present in *Nog* cKO HFs (Fig. 4c). These results suggest that epithelial deletion of *Nog* compromises but does not abolish SG fate specification and development. Furthermore, we also generated and examined *Foxc1* cKO (*Krt14-Cre/Foxc1^fl/fl^)* for SG development because FOXC1 was significantly reduced in miR-200 induced skin (Extended Data Fig. 3e), and FOXC1 regulates BMP production in HF-SCs^41^. However, although *Foxc1* cKO showed mild SG defects in adult (not shown) they did not show any discernible defects in SG morphogenesis by P5 (Fig. 4d-e). Taken together, these data suggest that perturbed BMP signaling pathway is partially responsible for compromised SG morphogenesis, and other mechanisms should mediate the blocked SG specification.

**Figure 4:**
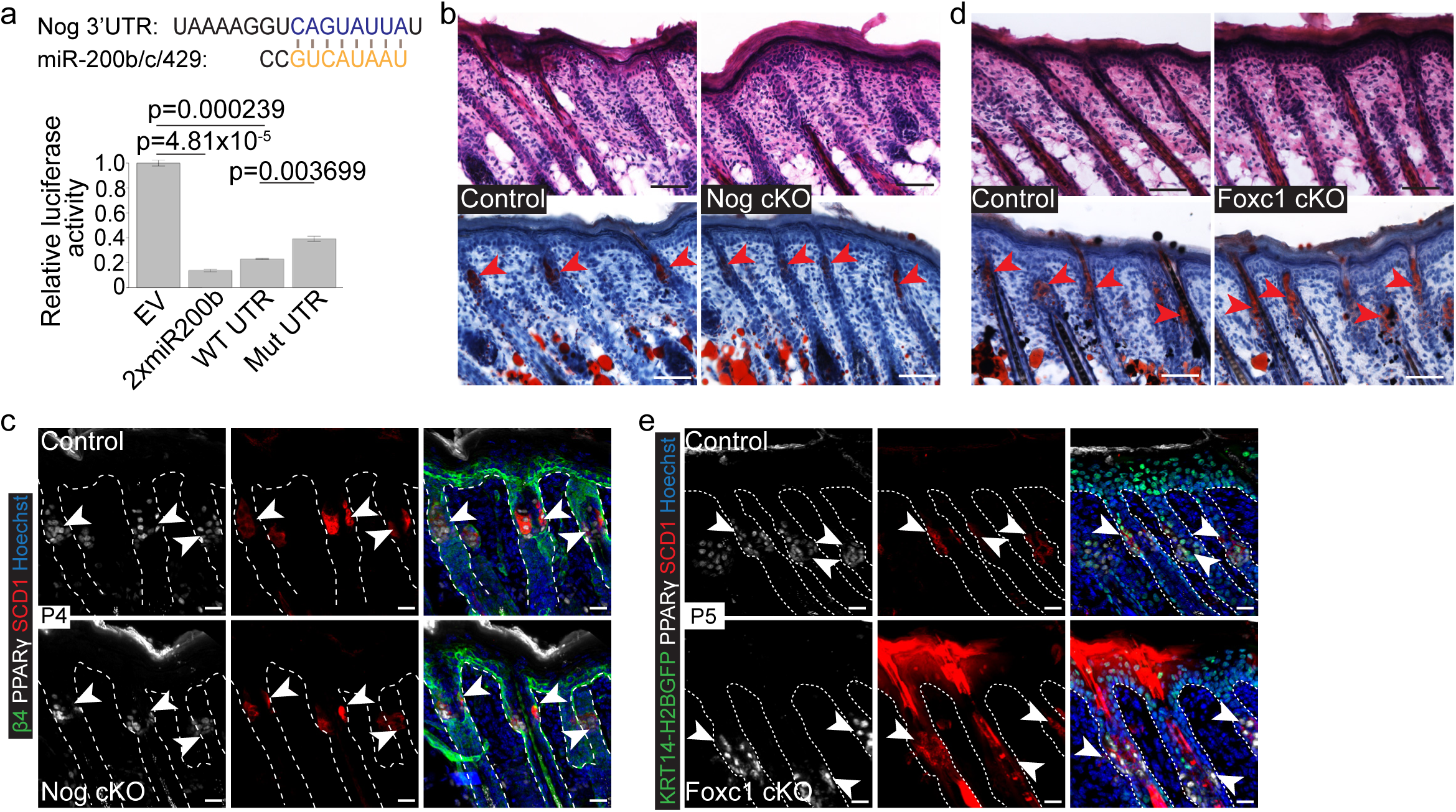
Perturbed BMP signaling does not abolish sebaceous gland formation. **a.** Schematics of *Nog* 3’UTR with the conserved miR-200b/c/429 target sequence. Luciferase reporter assay demonstrates the seed-dependent inhibition of *Nog* 3’UTR by miR-200b/c/429. The empty vector (EV) and a vector with 2x perfect matched miR-200b target sequences serve as negative and positive controls, respectively. Student’s t-test was performed for statistical analysis. **b.** H&E and Oil Red O staining demonstrates normal epidermal, HF development in *Nog* cKO skin **(H&E, top)** with minor SG maturation defect **(ORO, bottom)**. Red arrowheads in the ORO-stained sections point towards the SG. **c.** PPARγ and SCD1 IF of control and *Nog* cKO skin samples at P4 shows a slight reduction in SCD1^+^-mature sebocytes but no difference in PPARγ^+^ sebocytes. White dotted line outlines the HFs; white arrowheads point toward the SG. Scale bar: 20 μm. **d.** H&E and Oil Red O staining demonstrates normal epidermal, HF development in *Foxc1* cKO skin **(H&E, top)** and normal SG development **(ORO, bottom)**. Red arrowheads in the ORO-stained sections point to the SG. **e.** PPARγ and SCD1 IF of control and *Foxc1* cKO skin samples at P5 shows no difference in SG progenitors or late differentiated sebocytes, respectively. tdTomato signal in the FOXC1 cKO animal is marked by *. White dotted line outlines the HFs; white arrowheads point to the SG. Scale bar: 20 μm.

**Figure 5:**
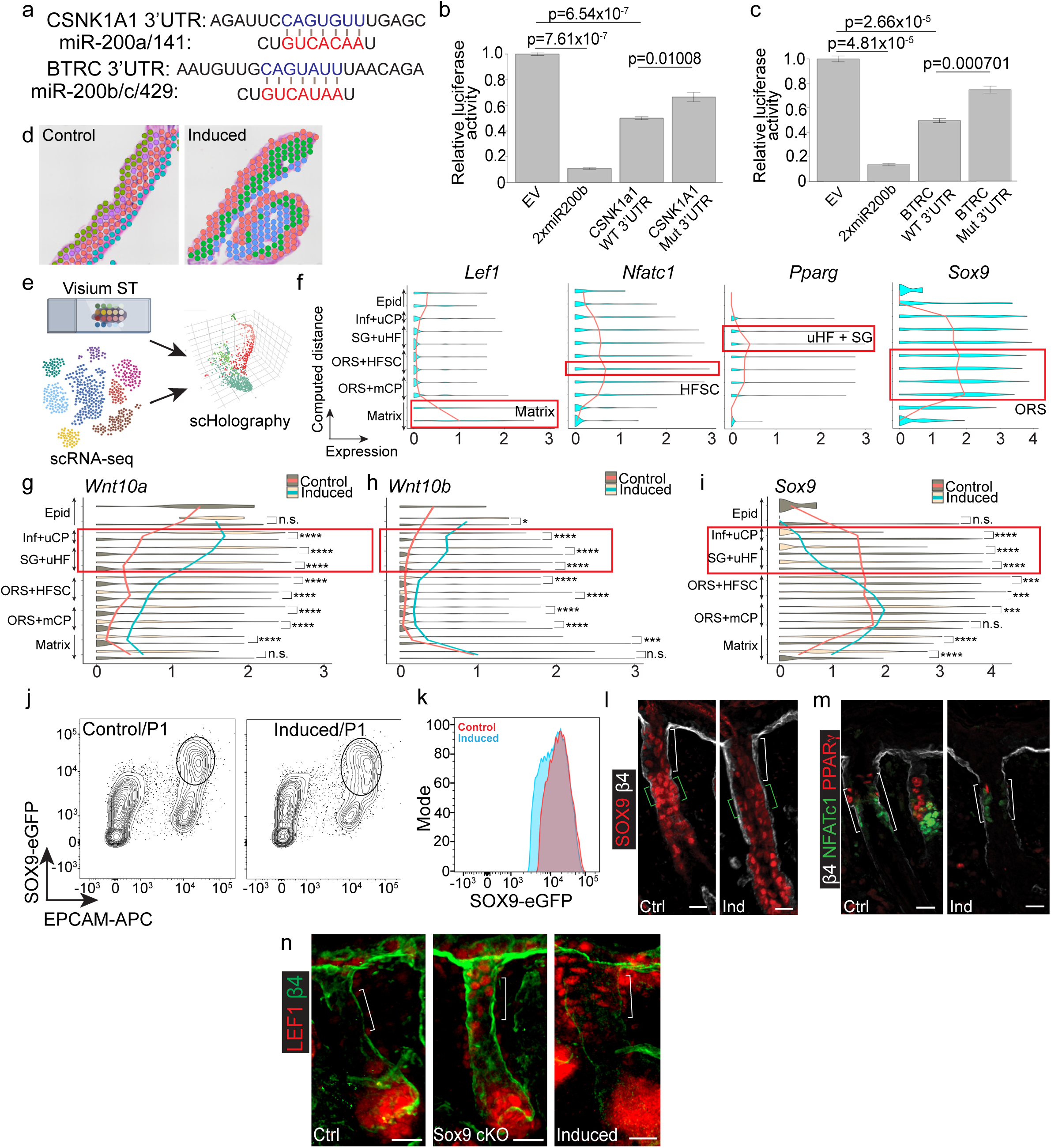
Spatial microenvironment analysis reveals compromised Sox9 expression in upper hair follicle. **a.** Conserved miR-200 targeting sequence on the 3’UTR sequences of *Csnk1a1* and *Btrc*. **b & c.** Luciferase reporter assay for *Csnk1a1* **(b)** & *Btrc* **(c)** showing the functional relevance of miR-200 binding sites to their respective 3’UTR. A Student’s t-test was performed for statistical analysis. **d.** Spatial transcriptomics clusters of control and induced skin at P3. **e.** Schematics of scHolography analysis from ST and scRNA-seq together. **f.** Expression of selected marker genes along the epidermal-to-matrix axis. The solid line connects the median gene expression level at each computed distance bin along the epidermal-to-matrix axis. **g-i.** Expression of *Wnt10a* **(g),** *Wnt10b* **(h)**, and *Sox9* **(i)** along the epidermal-to-matrix axis shows upregulation of *Wnt10a* and *Wnt10b*, as well as downregulation of *Sox9* in the upper HF region. Solid lines connect the mean expression level for individual genes for each sample across the spatial bins. Wilcoxon test was performed for statistical analysis. **j.** Representative contour plot for HF-keratinocytes marked by EpCAM-APC^+^/SOX9-eGFP^+^ from 2 control and 3 induced skin samples harvested at P1. **k.** Modal distribution of GFP intensity of the EpCAM^+^/SOX9-eGFP^+^ of both control (red) and induced (blue) cells. **l.** SOX9 IF shows uHF-specific downregulation in induced HF compared to control. Scale bar: 20 μm. **m.** NFATc1 and PPARγ downregulation in induced HF reveals compromised SOX9 function. Scale bar: 20 μm. **n.** In SOX9 cKO and miR-200 induced samples, LEF1+ cells were detected in the uHF region marked by a white bracket whereas these LEF1+ cells were absent in the control uHF. White brackets annotate the uHF region. Scale bar: 20 μm.

### Spatial microenvironment analysis reveals compromised Sox9 expression in upper hair follicle

Sebaceous gland is highly sensitive to the perturbation of WNT signaling^17,28,33,42^. Specifically, dominant negative LEF1 mutation, which compromises WNT signaling, leads to SG hyperplasia whereas activated WNT signaling through constitutively activated β-Catenin promotes ectopic hair formation at the expense of SG formation and maintenance^17,33^. We then turned to WNT signaling, whose negative regulators, *Csnk1a1* and *Btrc*, were identified as conserved miR-200 targets with the perfect 7mer sequences matching to the seed region (nucleotide No. 2-8) of miR-200s (Fig. 5a). We next validated them as *bona fide* miR-200 targets in the heterologous 3’UTR luciferase reporter assays (Fig. 5b-c). Inhibition of these negative regulators of WNT signaling by miR-200 was predicted to derepress WNT in upper HF regions, in line with elevated expression of *Lef1, Wnt10b,* and *Wnt10a* in miR-200 induced skin (Fig. 3i-j and Extended Data Fig. 3d).

To gain insights into the spatial gene expression pattern and microenvironment in developing skin, we applied 10x Visium ST platform to control and induced skin samples isolated from P3 mice (Fig. 5d). Upon QC and data processing (see Method), we detected 4 spatially distinct regions, including the epidermis, upper HF, lower HF and muscle, based on transcriptomic clustering of the spatial data in control (Extended Data Fig. 4a) whereas we only detected 3 distinct regions, including the epidermis, lower HF and muscle, in induced sample (Extended Data Fig. 4b). These results corroborated with our phenotypical and scRNAseq studies that upper HF region, marked by *Sfrp2* and *Krt79*, was conspicuously compromised. To further enhance the spatial resolution, we used our recently developed scHolography to reconstruct single-cell spatial neighborhoods by integrating scRNAseq and ST data with a machine-learning framework^43^ (Fig. 5e and Extended Data Fig. 5a). To confirm that scHolography spatially reconstructed skin appendages and enhanced quantitative analysis of spatial neighborhood, we computed the relative distance of major epithelial cell populations along the epidermal-to-matrix axis in control samples. Consistent with stereotypical skin and skin appendage organization, IFE cells were furthest away from the matrix cells, followed by infundibulum (Inf), uHF, junctional zone (JZ), SG, HF-SCs and ORS (Extended Data Fig. 5b-c). Furthermore, scHolography calculated the composition of epithelial cell clusters within spatially defined microenvironment, which allows spatial visualization of gene expression patterns^43^ (Extended Data Fig. 5d-e). We then plotted key marker genes associated with these distinct HF and SG populations, such as *Lef1*, *Nfatc1*, *Pparg* and *Sox9*, along the reconstructed spatial axis (Fig. 5f). Consistent with the well-established paradigm, *Lef1* expression was largely restricted to the Matrix region whereas *Sox9* expression was more broadly detected in upper HF regions and ORS but strongly repressed in the Matrix. Thus, the expression of *Lef1* and *Sox9* was mutually exclusive in developing HFs, reminiscent with their antagonistic patterns observed during hair placode formation^9^. As expected, the expression of *Pparg* and *Nfatc1,* markers of SG and HF-SCs, overlapped with *Sox9* in the uHF region. Interestingly, scHolography reconstruction correctly located *Pparg+* cells as more distal than *Nfatc1+* cells to the matrix, highlighting the enhanced spatial resolution and validating the computationally generated spatial neighborhoods achieved by the machine-learning framework.

The spatial map from both control and induced samples allowed the visualization and comparison of gene expression along the defined spatial axis. Indeed, when we visualized the expression patterns of *Wnt10a* and *Wnt10b,* we not only recapitulated their increased expression in induced samples but also pinpointed the cells from uHF regions as the most strongly elevated (Fig. 5g-h). Remarkably, we detected spatially confined reduction of *Sox9* in the same uHF regions where *Wnt10a/b* were strongly elevated (Fig. 5i). These results indicated that elevated WNT ligand expression was correlated with reduced *Sox9* expression in the uHF, reminiscent with the observation that WNT and SOX9 antagonize each other in the hair placode^9,44^. To confirm miR-200-mediated downregulation of *Sox9* expression, we bred miR-200 induced mice with *Sox9^IRES-EGFP^* knockin mice, which reports the transcriptional activity of the *Sox9* allele. In contrast to control, which showed uniformly strong GFP signals in neonatal skin, we observed not only the Sox9-GFP^hi^ population but also a Sox9-GFP^mid-lo^ population in induced skin (Fig. 5j-k and Extended Data Fig. 5f), This result suggested that *Sox9* transcription was downregulated in some appendage cells upon miR-200 induction, consistent with the spatial gene expression analysis (Fig. 5i).

These data prompted us to examine SOX9 expression by IF staining. Consistent with the spatial analysis of mRNA expression, we detected the reduction of SOX9 in uHF regions of the induced skin (Fig. 5l). Notably, NFATC1 signals were also reduced in *miR-200* induced skin, in addition to largely abolished PPARγ expression (Fig. 5m). When we knocked out *Sox9* by generating *Krt14-Cre/Sox9^fl/fl^*cKO, we observed that both NFATC1 and PPARγ expression was abolished (Extended Data Fig. 5g-h), demonstrating the requirement of SOX9 for the expression of these key TFs for SG and HF-SC progenitors, respectively. Interestingly, LEF1, which reflects *bona fide* WNT activities, gained ectopic expression in the uHF regions of both *Sox9* cKO and *miR-200* induced skin (Fig. 5n), revealing the antagonism of WNT and SOX9 in these progenitors. Taken together, these data reveal compromised SOX9 function in *miR-200* induced skin.

### SOX9 governs sebaceous gland gene expression program

Similar to miR-200 induced skin, hair morphogenesis of *Sox9* cKO skin proceeded whereas the SG formation was abolished^22^, revealing a differential requirement of SOX9 for the initial morphogenesis of skin appendages. Although SOX9 has been extensively studied for its role in maintaining HF-SCs^22,45–48^, how does SOX9 control SG-specific gene expression remain poorly understood. To investigate how SOX9 specifies the SG fate and how the process was disrupted by miR-200, we re-analyzed our P3 control scRNAseq data focusing on the uHF-to-SG trajectories. To identify genes that are critically involved in SG specification, we applied GeneTrajectory^49^ to deconvolve key gene programs among these uHF populations. Interestingly, GeneTrajectory analysis captured the uHF progenitor-to-SG transition, the cycling cell trajectory and uHF progenitor-to-uHF basal transitions as our cell trajectory analysis (Fig. 6a, Extended Data Fig. 6a-b and Table S2). Probing more deeply into the SG trajectory, which contributed primarily to the SG formation, we identified lipid and fatty acid metabolism as the most highly enriched GO terms among these SG trajectory genes (Fig. 6b). As expected, the cell cycle trajectory was highly enriched for genes associated with cell division and mitosis whereas the uHF trajectory was highly enriched for genes associated with keratinocyte differentiation and skin development (Extended Data Fig. 6c-d). These data collectively defined that the activation of lipid metabolism genes is the most prominent gene program during SG fate specification.

**Figure 6:**
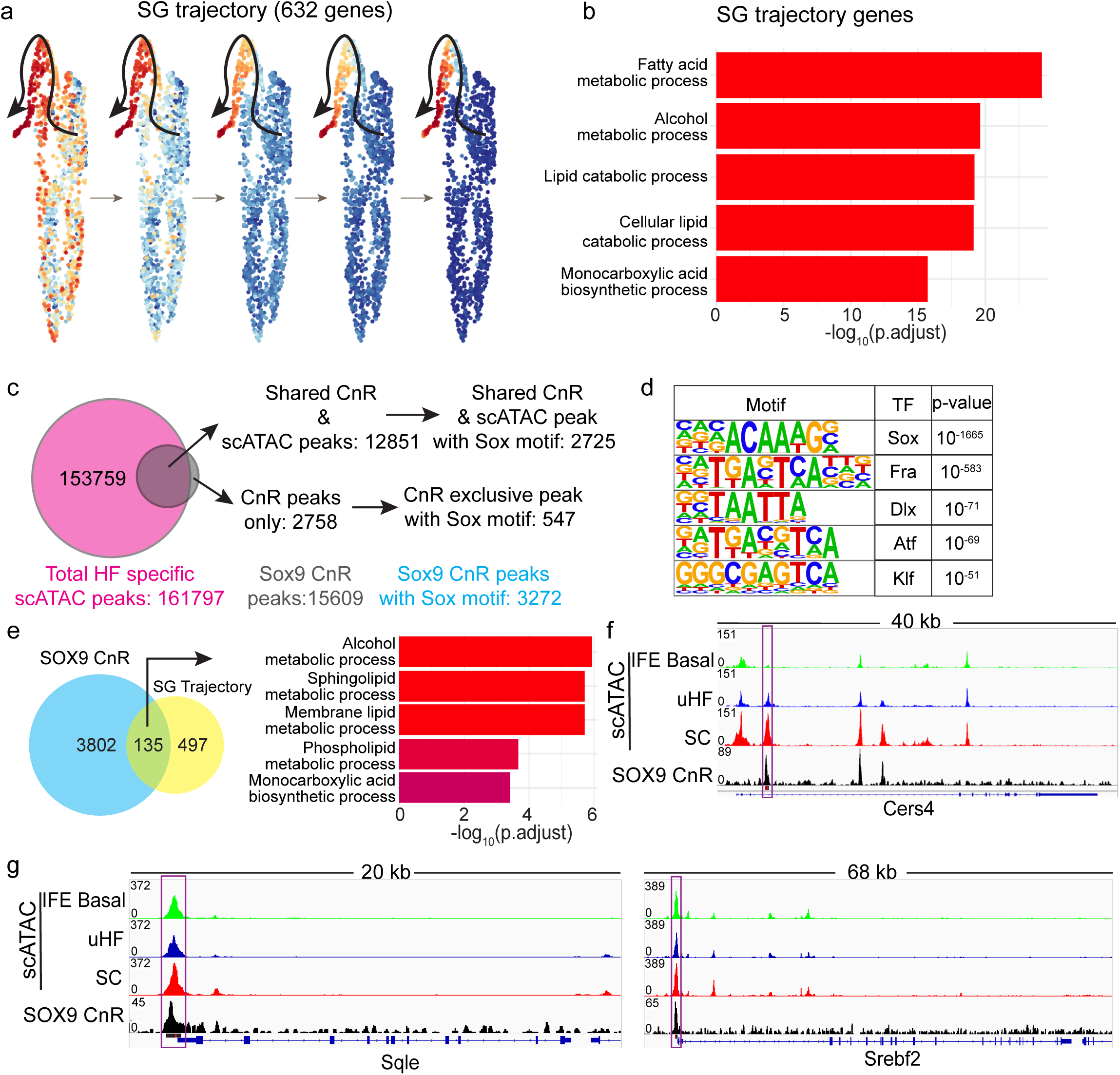
SOX9 governs sebaceous gland gene expression program. **a.** GeneTrajectory analysis of uHF and SG cells from control reveals a distinct trajectory from the uHF progenitors driving the SG fate. **b**. GO analysis of 632 genes from the SG trajectory. The top enriched terms are shown. **c.** Quantitative analysis of the peaks identified by SOX9 Cut&Run. **d.** The most highly enriched motifs (HOMER *de novo* motif search) in 3,272 SOX9 Cut&Run peaks. **e.** Intersection of genes identified in the SG trajectory with genes associated with SOX-motif containing peaks (left) and GO-term analysis of these SOX9-regulated SG trajectory genes (right). **f & g.** IGV tracks of representative genes Cers4 **(f)** Sqle **(g, left)** and Srebf2 **(g, right),** showing strong SOX9 Cut&Run signals on the promoter regions of these lipid metabolism genes. Purple rectangle annotates the peaks containing SOX motifs, a black horizontal line below the SOX9 Cut&Run track marks the actual peak with red vertical line pointing to the canonical ACAAAG motif for each peak.

To determine how SOX9 plays a role in SG fate specification, we performed SOX9 Cut n Run analysis by using sorted HF cells (Fig. 6c). Among 15,609 SOX9 Cut n Run peaks, 12,851 peaks were within open chromatin regions of HFs, detected by scATAC analysis (Extended Data Fig. 7). A total of 3,272 peaks contained the canonical SOX motif (Fig. 6d), suggesting direct binding of SOX9 to these DNA regions. By using the GREAT^50^ algorithm to assign individual enhancers and promoters to their regulated genes, we identified 3,937 genes as potential direct targets of SOX9. Interestingly, we identified 135 genes (21%) out of the 632 SG trajectory genes, including *Srebf2, Cers4*, *Lpin1* and *Scd2/3*, as SOX9 targets, and these genes function prominently in lipid and sphingolipid metabolism (Fig. 6e). Consistent with these global analyses, inspection of individual genes demonstrated strong SOX9 Cut n Run peaks containing SOX motif in gene locus, such as *Cers4, Sqle* and *Srebf2* (Fig. 6f-g). Taken together, these data reveal that SOX9 activates lipid and fat metabolism gene expression, which is characteristic of SG fate, in the uHF progenitor-to-SG trajectory.

### Mechanism of miR-200-mediated inhibition of sebaceous gland specification

Having established the key role of SOX9 in regulating SG gene trajectories, we next investigated how miR-200 induction blocks SG fate specification from uHF progenitors. Although both the specified SG cell cluster and cell cycling cluster were largely absent in miR-200 induced dataset, which prevented the analysis of miR-200’s role during the uHF progenitor-to-SG transition in our single-cell dataset, we reasoned that uHF progenitors, which harbor the progenitors to the SG^15^ and give rise to SG cells, should allow us to determine how induced miR-200 compromises SG related genes. Indeed, downregulated genes in induced uHF progenitors were highly enriched for genes associated with cell cycle, cell division and lipid metabolism. Interestingly, genes associated with electron transport, which reflects mitochondrial functions, were also downregulated (Fig. 7a). These data suggested that induced *miR-200* expression compromises the cell cycle progression and the activation of lipid metabolism genes, which are necessary for the SG fate specification. By overlapping CLEAR-CLIP identified miR-200 targets and downregulated genes detected in these uHF progenitors, we identified 136 miR-200 targets that were downregulated in the progenitors (Fig. 7b). GO term analysis revealed that these genes were predominantly enriched for genes involved in cell cycle and cell division, lending further support to the potent role of miR-200s in inhibiting cell proliferation (Fig. 7b-c). Many of these genes, including *Ccng2, Ccnd1, Mcm4* and *Egfr*, were experimentally validated miR-200 targets^6,7^. Interestingly, 70 of the SG trajectory genes were also downregulated in these uHF progenitors, and they were highly enriched for SOX9-targeted lipid metabolism genes, including *Sqle* (Fig. 7d). Thus, the effect of compromised SOX9 function was already evident in these progenitors, providing a molecular explanation for the observation that miR-200 induced uHF progenitors were unable to give rise to the SG. Taken together, these results demonstrated that miR-200-mediated inhibition of cell cycle progression together with compromised SOX9 function, which is required for the activation of lipid metabolism genes, blocks the SG fate specification (Fig. 7e).

**Figure 7:**
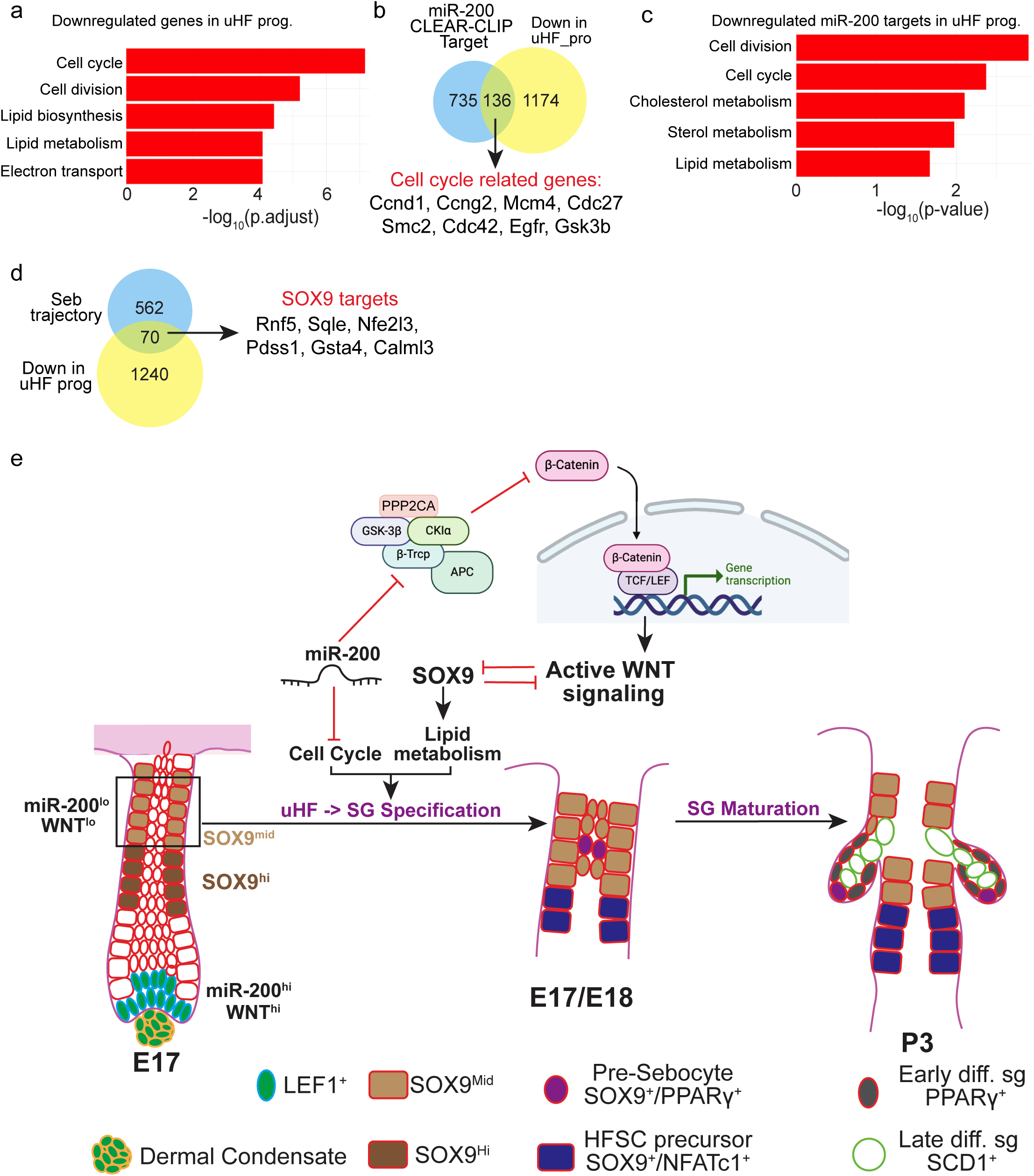
Mechanism of miR-200-mediated inhibition of sebaceous gland specification. **a.** GO-terms of the downregulated genes in uHF progenitors upon miR-200 induction are enriched for genes associated with cell cycle and lipid metabolism. **b.** 136 of the CLEAR-CLIP identified miR-200 targets are downregulated in the uHF progenitors, as shown by the Venn diagram. Selected cell cycle genes that are miR-200 targets are listed. **c.** GO term analysis of miR-200 target genes that are downregulated in the uHF progenitors. **d.** Intersection of the SG trajectory genes with downregulated genes in uHF progenitors reveals the enrichment for SOX9 targets. **e.** Schematics of miR-200 mediated inhibition of sebaceous gland specification.

## Discussion

In this study, we leverage the highly choreographed morphogenesis of skin appendages as a precision model to investigate how miR-200s regulate epithelial plasticity. We demonstrate that the physiological exclusion of miR-200s from the upper HF is required for sebaceous gland specification. Our previous work established that miR-200s are enriched within the early hair germ, overlapping with LEF1+/WNT^hi^ progenitors during placode formation^7^. Here, by using miRNA FISH to examine more mature postnatal HFs, we show that this spatial restriction to WNT^hi^ cells is maintained such that miR-200 expression remains confined to the WNT^hi^ hair matrix and is conspicuously absent from the upper HF - the specific niche where SGs and HF-SCs arise from a WNT^lo^ microenvironment. Using an inducible mouse model, we find that forced expression of miR-200s in the WNT^lo^ uHF region potently blocks SG specification while surprisingly sparing HF morphogenesis. Thus, miR-200s expression appears to track with WNT^hi^ cells, and its induction in uHF acts as a strong blockade to the SG fate, revealing an unexpected specificity of miR-200s in governing epithelial plasticity.

To determine the molecular basis of this blockade, we combine genome-wide target capture (CLEAR-CLIP), which identifies physical miRNA-mRNA interactions within a defined cellular context^6^, with Targetscan prediction, which leverages evolutionary conservation and local sequence context^40^. This integrated approach identifies and validates components of the β-catenin destruction complex, specifically *Csnk1a1* and *Btrc*, as miR-200 targets. These findings suggest a model where miR-200 expression reinforces the WNT^hi^ state in both hair germ and hair matrix. Conversely, in the uHF where WNT signaling must be attenuated to permit SG specification, ectopic miR-200 expression reactivates WNT signaling, leading to the repression of SOX9. Although WNT perturbation has long been known to interfere with SG development^17,42^, its link to SOX9, the master regulator of the HF lineages^22,45,47^, has not been fully established in this context. Leveraging our recently developed scHolography framework^43^, we significantly improve the spatial resolution of 10x Visium spatial transcriptomics to pinpoint this antagonism in a spatially defined manner, revealing that derepressed WNT signaling spatially correlates with compromised SOX9 function in the uHF upon miR-200 induction. Notably, SOX9 is known to dampen WNT activity by recruiting the destruction complex and competing with TCF/LEF for nuclear β-catenin^51,52^. We therefore propose that miR-200-mediated WNT activation and the subsequent downregulation of SOX9 form a robust feedforward loop that strongly compromises SOX9 function. This defines a critical WNT-SOX9 antagonism in the uHF that is spatiotemporally distinct from their well-appreciated antagonism in the hair placode^9,44^.

Finally, we elucidate the mechanism by which SOX9 specifies the SG fate. While genetic deletion of *Sox9* is known to cause a complete failure of SG formation^22^, the downstream effectors remains elusive. Combining SOX9 Cut&Run and GeneTrajectory analysis, we identify a direct regulatory axis where SOX9 binds to high-affinity motifs in the promoters and enhancers of a broad lipid metabolic network, including the master regulator *Srebf2*, the ceramide synthase *Cers4* and the rate-limiting enzyme *Sqle*. This finding expands the role of SOX9 from a lineage specifier to a metabolic licensor. We propose that SOX9 expression not only turn on skin appendage genes but also activate the high-output biosynthetic capacity required for lipid biogenesis and sebum production in the upper HF. Furthermore, our GeneTrajectory analysis reveals that cell cycle progression, a process potently inhibited by miR-200s^1,7^, is intimately linked to this metabolic programming during SG specification. Thus, the coordinated inhibition of the SOX9-dependent lipogenic axis and key cell cycle drivers, mediated by miR-200s, blocks the SG formation. In a broader context, our findings provide a new perspective on the tumor suppressor function of the miR-200 family. Because SOX9 plays critical roles for both skin appendage formation^22,45^ and self-renewal of tumor initiating cells of basal cell carcinoma^26^, we propose that this miR-200-mediated SOX9 repression represents a common mechanism that restricts epithelial plasticity during development and underlies the tumor suppressor functions of this miRNA family.

## Methods

### Mice

All experiments were carried out following IACUC-approved protocols and guidelines at Northwestern University. Mice were bred and housed according to the IACUC guidelines in a pathogen-free facility at the Feinberg School of Medicine, Northwestern University. Housing rooms have 12-hour light/dark cycles with an ambient temperature of 23°C and 50% humidity.

The Tg mouse was generated as described previously^7^. Pregnant female mice were fed with Doxycycline chow at 625 mg/kg (Inotiv rodent diet TD.01306) to induce the expression of miR-200b cluster miRNAs from E10.5 (unless mentioned otherwise) until sample collection.

The Nog^flox/flox^ mouse was a gift from Dr. J. Kessler, Northwestern University. The Foxc1^flox/flox^ mouse was a gift from Dr. T. Kume, Northwestern University. SOX9^flox/flox^ (B6.129S7-*Sox9^tm2Crm^*/J) mouse was obtained from the Jackson Laboratory (strain number: 013106). All these three mouse strains were bred with Krt14-Cre or Krt14-H2BGFP animals. SOX9^IRES-eGFP^ (Tg(Sox9–EGFP)EB209Gsat) mouse was obtained from MMRRC (stock number: 011019-UCD) and was bred with *Krt14-rtTA/pTRE2-miR-200b/a/429* mouse.

### In situ hybridization

In situ hybridization for miR-200b was done following the protocols established previously^53,54^ with minor modifications. We treated our samples with 4µg/ml proteinase K instead of 20µg/ml. The samples were hybridized with a double-digitonin locked probe against miR-200b at 51°C for 2 hours. Sections were incubated with β4-integrin antibody (1:100; clone 346-11A BD Pharmingen #553745) and PPARγ antibody (1:200; clone 81B8; Cell Signaling Technology #2443) together for 2 hours at RT, followed by DyLight 549 and Alexa Fluor 647 conjugated secondary antibodies against rat and rabbit, respectively, for 1 hour at RT.

### Histological staining

H&E and Oil Red O (ORO) stainings were performed at the Pathology Core at Robert H Lurie Comprehensive Cancer Center, Northwestern University. Briefly, OCT-embedded dorsal skin samples were sectioned to 12 μm thickness. For H&E staining, sections were fixed in 95% alcohol for 10 mins at RT, then incubated with hematoxylin and eosin for 7 mins and 3 mins respectively with frequent washes. The sections went through a series of dehydration steps and were finally mounted with Permount mounting medium. For ORO staining, sections were briefly washed with DI water and incubated with 60% ORO for 10 mins at RT. The extra stain was removed, and sections were washed in DI water. Nuclei were counterstained with Harris Hematoxylin (Surgipath 01560) for 1 min at RT, followed by a wash with DI water. Sections were blued using 0.4% ammonia water, and washed well in DI water. Finally, the section was mounted using an aqueous mounting media. Both H&E and ORO images were acquired using a Leica DM5500B microscope with a Leica camera and Leica LAS X suite v3.7 software.

### Immunofluorescence

OCT-embedded dorsal skin samples or embryos were sectioned to 25μm. Sections were fixed in 4%PFA for 15 mins at room temperature (RT), and permeabilized with 0.3% Triton X-100 for 10 mins at RT. Sections were blocked with 5% normal serum of the same species in which the secondary antibody was raised and 0.1% Triton X-100 for 1 hour at RT before overnight incubation at 4°C with primary antibodies diluted in blocking buffer. For antibodies raised in mouse, MOM mouse Ig blocking reagent was used (Vector Labs #BMK-2202) to block for 1 hour at RT, followed by two brief PBS washes at RT. Then the samples were incubated in MOM protein concentrate for 5 mins at RT and the samples were incubated with primary antibody diluted in the MOM protein concentrate overnight at 4°C. The next morning, the antibody cocktails were removed, and sections were washed three times in PBS for 5 mins each at RT. Sections were incubated with secondary antibodies conjugated with either Alexa Fluor 488/555/647 (at 1:2000 dilution) at RT for 2 hours. Nuclei were counterstained with Hoechst 33342 (Invitrogen) at 1:3000 dilution. Slides were mounted with Fluoromount G (Southern Biotech). Imaging was performed on a Nikon A1 confocal microscope and viewed using Imaris software. Primary antibodies and dilutions used in this study are: PPARγ (1:400; Cell Signaling Technology #2443), SCD1 (1:400; Santa Cruz Biotechnology #sc-14719), LEF1 (1:400; Cell Signaling Technology #2230), FOXC1 (1:100; Cell Signaling Technology #8758), KRT5 (1:2000; Covance #SIG-3475), β4 integrin (1:400; clone 346-11A; BD Pharmingen #553745), SOX9 (1:200; Milipore Sigma #HPA001758), Alexa Fluor 488 conjugated with SOX9 (1:200; Millipore Sigma #AB5535-AF488), NFATc1 (1:10; Developmental Studies Hybridoma Bank #7A6). QKI antibody (1:500; Bethyl Laboratories #IHC-00574) was a gift from Dr. Jian Hu at MD Anderson Cancer Center.

### Single-cell RNA-sequencing library preparation

Total dorsal skin from one pair of P3 control and Tg animals were surgically removed. The epidermis and dermis were separated after incubating the dorsal skin with 2.5U/mL Dispase for 1 hour at 37°C. The epidermal layer was treated with 0.05% Trypsin-EDTA for 8 minutes at 37°C, and a single-cell suspension was prepared. The dermal layer was mined in smaller pieces and treated with 1X Collagenase IV (Worthington, LS004188) for 90 minutes at 37°C. We enriched for HFs by transferring the dissociated dermal layer in a tube and kept it on ice for 3 minutes. Intact HFs that precipitated at the bottom of the tube were treated with 0.05% Trypsin-EDTA for 8 minutes at 37°C and passed through a 40 μm filter. Cells were pelleted down. The epidermal and HF-containing fragments were mixed at a 30:70 ratio to enrich for epithelial cells. Finally, the libraries were constructed following 10x genomics Chromium single cell 3’ reagent kits v3.1 (PN-1000120). The libraries were sequenced on the Illumina NovaSeq 6000 platform to achieve an average of approximately 50,000 reads per cell.

### scRNA-seq analysis

Cellranger Single-Cell Software Suite (v6.0.0, 10x genomics) was used to perform barcode processing, mapping to mm10 reference genome and single-cell 3’ gene counting. Barcodes, features, and matrix files were loaded into SoupX 1.6.2^55^ to remove ambient RNA contamination. These SoupX-corrected files were used to find doublets using DoubletFinder 2.0.4^56^, and only barcodes associated with singlets were exported. We loaded SoupX-corrected barcodes, features, and matrix into Seurat 4.3.0 for downstream analysis. Low-quality and dead cells were excluded by using filtering criteria nFeature_RNA (>1000 and <6000) and mitochondrial percentage (<20%). We regressed out any variation because of genes associated with the cell cycle using Seurat *CellCycleScoring* and *SCTransform* algorithm. After UMAP dimensional reduction and clustering, we identified clusters using known genes (Extended Figures 1 and 3). To analyze epithelial cells in higher resolution, we subset epithelial cells and re-ran the analysis pipeline. We further subset upper HF and sebocytes for a higher resolution transcriptomic view and pseudotime analysis using scVelo 0.3^38^. Gene trajectory analysis was performed using GeneTrajectory1.0.0^49^.

For cluster-specific differential gene expression analysis, we exported the barcodes of each cell, and they were randomly split into two groups to make pseudo-replicates. We parsed the BAM file according to the barcodes, and the expression of each gene for the cells in each pseudo-replicate was calculated from the aligned BAM file by HTSeq-count. Differential gene expression analysis was performed using the DESeq2 package in R^57^. Gene ontology analysis for up- or downregulated genes was done using ClusterProfiler4.0^58^ on the GO_BP, GO_CC, GO_MF, Hallmark and WP database from EnrichR^59^ and using UP_KW_Biological process on DAVID Bioinformatics^60^.

### Spatial transcriptomics (ST) library preparation and data analysis

ST was performed on control and Tg dorsal skin samples harvested at P3. Harvested dorsal skins were fixed in 4% PFA overnight and embedded in paraffin. Prior to ST, RNA was extracted from 10 μm thick sections for each FFPE block using Qiagen RNeasy FFPE Kit (73504). The RNA quality was assessed by calculating the DV200 value using an Agilent 2100 bioanalyzer.

For ST, 5 μm thick section from the FFPE block was sliced using a microtome and placed on the capture areas on a 10x genomics Visium Spatial Gene Expression Slide (PN-2000233). Deparaffinization, H&E staining following recommendations from 10x genomics (CG000409). Images for H&E-stained sections were acquired using a Nikon Ti2 widefield microscope. Visium Spatial Gene Expression libraries were constructed using the 10x genomics FFPE reagent kit (PN-1000361), mouse transcriptome probe kit (PN-1000365) following the protocol from 10x genomics (CG000407). The libraries were sequenced on the Illumina NovaSeq 6000 platform.

The sequencing files were first aligned to mouse reference genome mm10 by Space Ranger v1.3.1. The Seurat v4.3 package was used for data processing and visualization. The ST data was integrated with stage-matched scRNA-seq data, and downstream analysis was performed using scHolography v0.1.

### Cell suspension preparation and Flow cytometry for *Sox9^IRES-EGFP^* analysis

Dorsal skin samples from control and induced pups at P1 were minced into smaller pieces using scalpel blades, and incubated with 1 ml of 1X collagenase IV for 1.5 hours at 37°C on a rocker. An equal amount of 1X HBSS was added to each sample, and digested skin tissues were pelleted down by centrifuging at 500g for 8 minutes at 4°C. Supernatant was removed, cell pellet was resuspended in 1 ml of 0.05% pre-warmed Trypsin-EDTA and incubated at 37°C for 8 minutes. Trypsin-EDTA was neutralized using 3% chelated FBS in 1X PBS, filtered cells using 40 μm strainers, and pelleted the cells using the same conditions as before. Cells were resuspended in 3% chelated FBS in 1X PBS after removing the supernatant and stained with EpCAM-APC (1:800; BioLegend 118213) for 1 hour on ice, in dark. After staining, cells were washed with 3% chelated FBS in 1X PBS twice. Finally, cells were resuspended in 1X PBS containing Hoechst 33342 (1:3000) and ran on a BD Fortessa flow cytometer. Flow cytometry data were analyzed using FlowJo.

### Sorting for CUT&RUN

SOX9 Cut-n-Run was performed on Krt14-H2BGFP^+^ hair follicle (HF) keratinocytes at the P3 stage. Epidermis from the dorsal skin was peeled off after treating the skin with 2.5U/mL Dispase at 37°C for 1 hour. Remaining dermal fragments harboring HFs were treated with Collagenase IV for 1.5 hours at 37°C. Digested samples were filtered through a 40 μm strainer and pelleted down at 500g for 5 mins. Cell pellet was treated with 0.05% Trypsin-EDTA for 8 minutes at 37 °C. Trypsin was neutralized by adding an equal amount of 1X PBS with 3% chelated FBS. Cells were pelleted down and resuspended in 3% chelated FBS in 1X PBS and 1:3000 Hoechst 33342. Finally, cells were filtered through a 50 μm filter, and Hoechst^lo^/*Krt14-H2BGFP*^+^ HF-keratinocytes were FACS-sorted using a MACSQuant Tyto instrument.

### CUT&RUN

SOX9 Cut-n-Run was performed on sorted Hoechst^lo^/*Krt14-H2BGFP*^+^ HF-keratinocytes at the P3 stage. Approximately 300,000 cells were used to prepare each Cut&Run library. Antibody binding, targeted chromatin release, and DNA purification were performed using Epicypher CUTANA kit (#14-1048) following manufacturer’s recommendation and previously published protocol by Law et al., 2024. The sequencing libraries were constructed using NEB Ultra^TM^ II DNA Library Prep Kit for Illumina (E7645). Anti-Sox9 antibody (Millipore Sigma, #AB5535) was used for the Cut&Run.

Paired-end Cut&Run reads were aligned to the mouse genome mm10 using Bowtie2. We merged the replicate BAM files and performed peak calling using MACS2. Cut-n-Run peaks were then annotated with the previously reported SOX10 motif using HOMER *annotatePeaks.pl* for downstream SOX9 target analysis.

### Single-cell ATACseq (scATAC) library generation

ScATAC library was prepared using a similar approach as scRNA-seq library. To enrich the epithelial cells, we mixed epidermal to dermal fragments at an 80:20 ratio after the respective fragments were isolated following the same method as scRNA-seq library preparation. The library was prepared using 10X genomics v2 kit.

### RT-qPCR

Control and induced epidermis were separated after incubating the respective dorsal skin in 2.5U/mL Dispase at 37°C for 1 hour. Separated epidermis samples were lysed in TRIzol (Ambion #15596018), and RNA was isolated using the manufacturer’s recommendation. cDNA was prepared from 500 ng of RNA for each samples using SuperScript III RT kit (ThermoFisher #11752050). Finally, qPCR was performed using gene-specific primers (Table S3) and iQ SYBR green (BioRad #1708880).

### Luciferase assay

3’UTRs of the respective genes were PCR amplified from cDNA using the primers listed in supplemental table S3. miR-200 binding site mutations were carried out by PCR using WT 3’UTR as the template DNA and primers harboring designated mutation sequence (Table S3). Both WT and mutant 3’UTRs were cloned in pGL3 vector (Promega). 5 ng of Renilla luciferase, 20 ng of pGL3 reporter and 380 ng of MSCV-IRES-GFP retroviral vector (MIGR) were co-transfected into mouse keratinocytes using the LT-1 transfection reagent (Mirus Bio #MIR2300). Cells were lysed after 48 hours post-transfection, and reporter assays were performed for both renilla and firefly luciferase using the Dual-GLO luciferase assay system (Promega #E1910).

### Statistical analysis

Statistical analyses were performed for all the indicated data using two-tailed Student’s t-test, except scRNA-seq and ST-related data quantification, where a Wilcoxon test was performed. Exact *P* values are mentioned for the two-tailed Student’s t-test. For the Wilcoxon test, statistical significance is indicated by asterisks (*: *P* < 0.05, **: *P* < 0.005, ***: *P <* 0.0005, ****: *P* < 0.0005) in the figures. Sample size (n) of each dataset is mentioned in the corresponding figure legends.

## Data availability

All data supporting the findings of this study are available within the paper and the supplementary files. Raw sequencing data and corresponding processed data related to this study have been deposited in the Gene Expression Omnibus under accession codes GSE312275 (SOX9 Cut&Run), GSE312276 (E18 control scATAC-seq), GSE312278 (P3 control and miR-200 induced scRNA-seq), and GSE312281 (P3 control and miR-200 induced Spatial Transcriptomics). Source data are provided with this paper. All other data supporting the findings of this study can be available from the corresponding author upon request.

## Code availability

All bioinformatic tools and pipelines used to analyze data related to this study are mentioned in the methods section. Codes can be available from the corresponding author upon request.

## Acknowledgements

We thank E. Fuchs (Rockefeller University, HHMI) for the Krt14-H2BGFP, Krt14-Cre, and Krt14-rtTA mice, J. Kessler (Northwestern University) for the Nog^fl/fl^ mice, K. Green, R. Lavker, F. Yue (Northwestern University) for their advice and comments. We also thank C. Arvanitis, M. De Niz, P. Dluhy at the Nikon CAM facility for their technical assistance, B. Frederick at Pathology Core Facility at RH Lurie Cancer Center for technical assistance, and S. Pandey at the Immune Assessment Core for single cell genomics. This study was supported by R01AR066703. R.Y. was supported by R01AR066703, R01HD107841, R01AR081103 and R01AR071435. We also acknowledge the support from NU-SBDRC (P30AR075049) and the Che Foundation.

## Author Contributions

R.Y. conceived the study. A.D., D.W., and R.Y. designed the experiments. A.D. carried out experiments and phenotypical characterizations with A.C. and M.A.W. A.D. performed data analysis with assistance from D.W. and Y.C.F. for the single-cell experiments. Y.C.F. performed single cell and spatial mapping and scHolography analysis. A.D. and H.L. prepared the CUT&RUN library for SOX9. D.W. and H.L. prepared the scATAC-seq library. G.B., A.S., and J.H. assisted sample preparations and tissue embedding at different stages of the project.

## Materials & Correspondence

Material requests and correspondence should be directed to R.Y.

**Extended Data Figure 1: Single-cell RNA sequencing analysis of neonatal skin. a.** Schematic representation of the scRNA-seq library preparation and data analysis pipeline. **b.** UMAP representation of the clusters identified in control dorsal skin at P3. A red dotted circle outlines the epithelial cells. **c-e.** FeaturePlot **(c,d)** and Dotplot **(e)** expression pattern of known genes to annotate epithelial **(c, red rectangle in e)** and non-epithelial clusters **(d, green rectangle in e)**. Red dotted circles in **(d)** annotate the individual cell population.

**Extended Data Figure 2: Spatiotemporal patterns of sebaceous gland specification and morphogenesis. a.** H&E staining of control and induced dorsal skin at P3. White dotted lines outline the sebaceous gland in control HF. Scale bar 20 μm. **b-d.** IF staining of PPARγ (green) and SCD1 (red) in different hair follicle developmental stages at E17 (**b**), E18 (**c**), and P1 (**d**) stages demonstrates the appearance of the early and late sebocytes. White: KRT5 (**b**) and β4-integrin in (**c, d**); scale bar: 20 μm. White arrowheads in the red inset in (**c)** highlight the appearance of basal PPARγ^lo^ cell, whereas the suprabasal cell is PPARγ^hi^.

**Extended Data Figure 3: Single-cell RNA sequencing analysis of upper hair follicle regions. a.** UMAP representation of only epithelial cells from Extended fig. 1b. **b.** DotPlot for various known genes was used to annotate the clusters in **(a)**. The blue rectangle highlights the SG cluster and associated gene expression. **c.** Violin Plots for key marker gene to identify uHF clusters. The red solid line separates the four uHF-related and two SG-related clusters. **d.** IF for LEF1 (green) at E18 with control and induced dorsal skin samples shows elevated LEF1 expression in the miR-200b/a/429 induced HF. Scale bar: 20 μm. **e.** IF for FOXC1 (red) at E18 with control and induced dorsal skin samples, shows reduced FOXC1 expression in induced HFs. Scale bar: 20 μm.

**Extended Data Fig. 4: Spatial transcriptomics analysis of control and miR-200 induced skin. a.** & **b.** Marker gene expression from each ST cluster, identified in control **(a)** and induced **(b)** dorsal skin at P3.

**Extended Data Fig. 5: Spatial transcriptomics analysis with scHolography reconstructs the hair follicle architecture and microenvironment. a.** Schematic representation of scHolography workflow. **b & c.** scHolography computed distance of each cluster with Matrix as reference cluster shows reliable reconstruction of control **(b)** and induced **(c)** HFs. Wilcoxon test was performed to check for statistical significance. **d & e.** Reconstructed HF architecture by plotting proportions of each HF-related single-cell clusters along the computed epidermal-matrix axis for control **(d)** and induced **(e)** HF. **f.** Quantification of the median GFP intensity of the EpCAM^+^/SOX9-eGFP^+^ cells from control (n=2) and induced (n=3) dorsal skin at P1. A t-test was performed to check statistical significance. **g.** IF for NFATc1 (red) shows absence of future HF-SCs in SOX9 cKO HFs at P3, whereas these cells are present in control HFs. The white bracket in control outlines the stem cell compartment missing in SOX9 cKO. Scale bar: 20 μm. **h.** IF for PPARγ (green) identifies lack of sebocytes in SOX9 cKO HFs, similar to miR-200 induced HFs at P3. A white bracket in control outlines the SG compartment missing in SOX9 cKO. Scale bar: 20 μm.

**Extended Data Fig. 6: GeneTrajectory analysis of upper hair follicle lineages. a. & b.** Two distinct trajectories, cell cycle and upper hair follicles, that drive the uHF fate. **c. & d.** GO-term analysis with genes from **(a)** & **(b)**, respectively, shows trajectory in **(a)** is enriched with cell cycle genes whereas trajectory in **(b)** is associated with differentiation genes.

**Extended Data Fig. 7: Single-cell ATAC-seq identifies distinct cell clusters based on their open chromatin landscape. a.** UMAP visualization of distinct clusters based on differential chromatin opening at E18. **b.** IGV-track representation of the clusters from **(a)**, for key genes to identify and annotate each cluster.

**Table S1: List of Epidermis vs Appendage genes (related to Fig. 1**).

**Table S2: List of genes from GeneTrajectory analysis (related to Fig. 6 and Extended Fig. 6).**

**Table S3: List of primers used in this study.**

