## Supplementary figures and images for "Coordinated inhibition of SOX9 and cell cycle progression by microRNA-200 restricts sebaceous gland fate specification"

### Extended Data Figure 1

##### Extended Data Figure 1

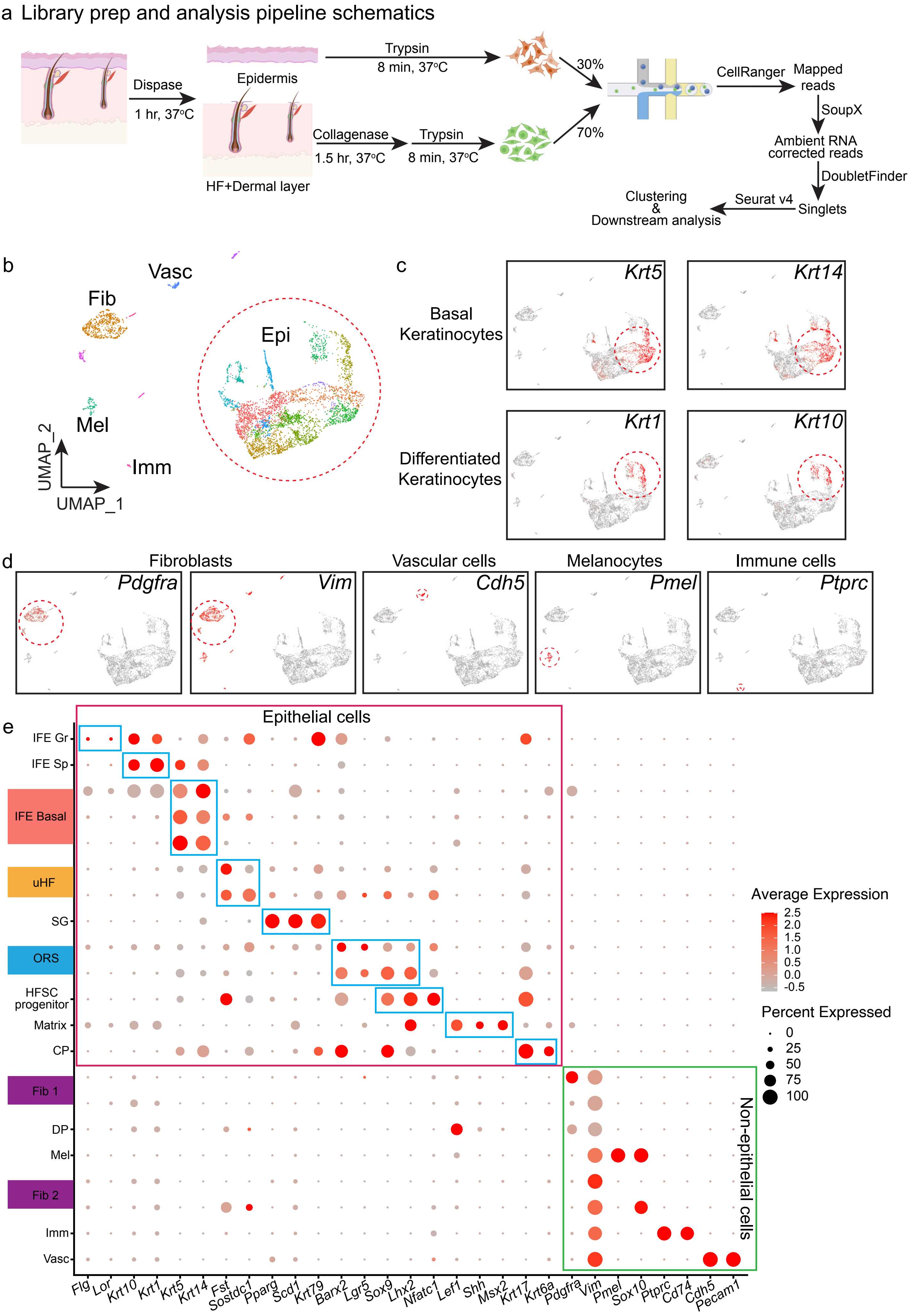

### Extended Data Figure 2

#### Extended Data Figure 2

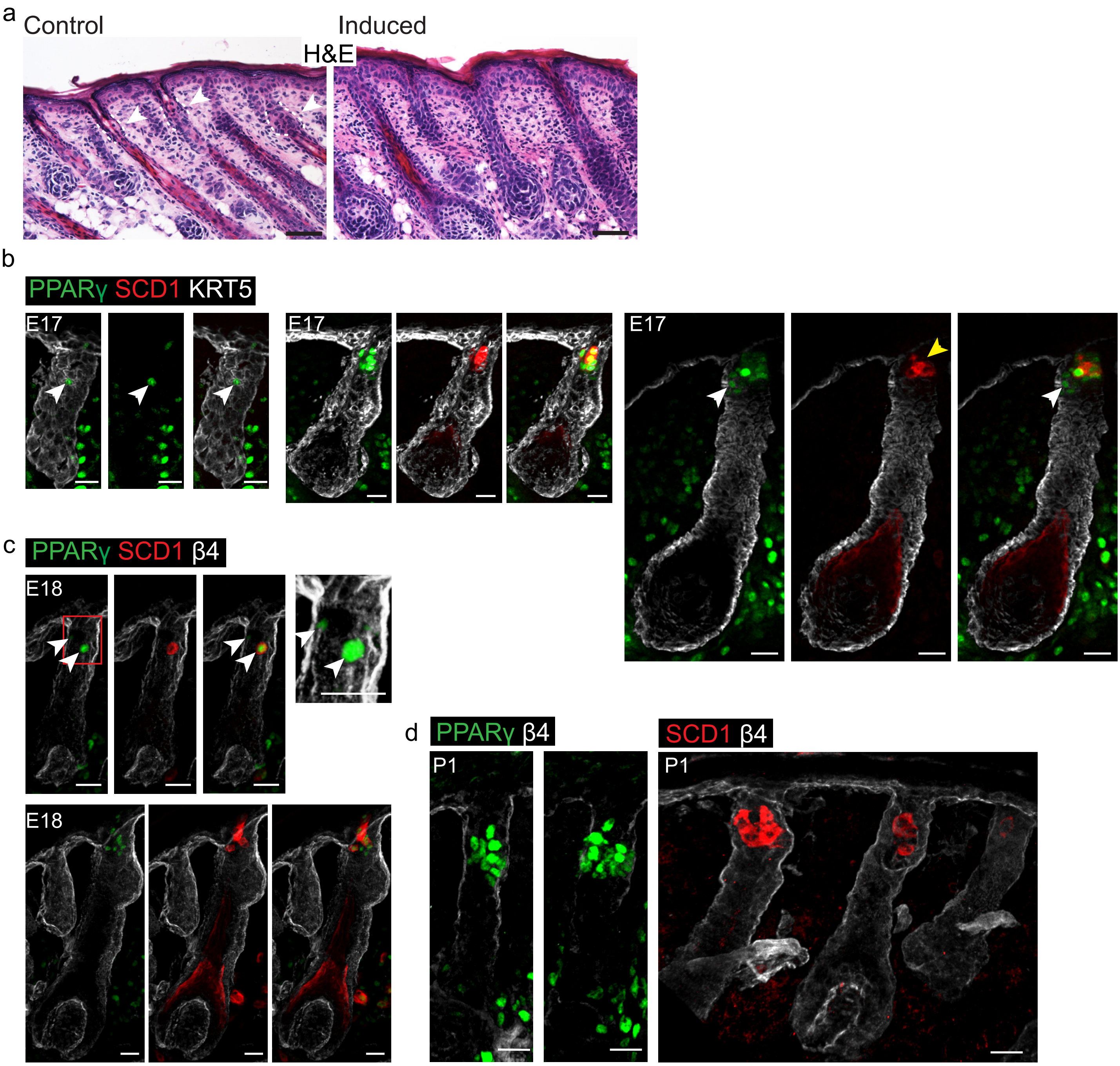

### Extended Data Figure 3

#### Extended Data Figure 3

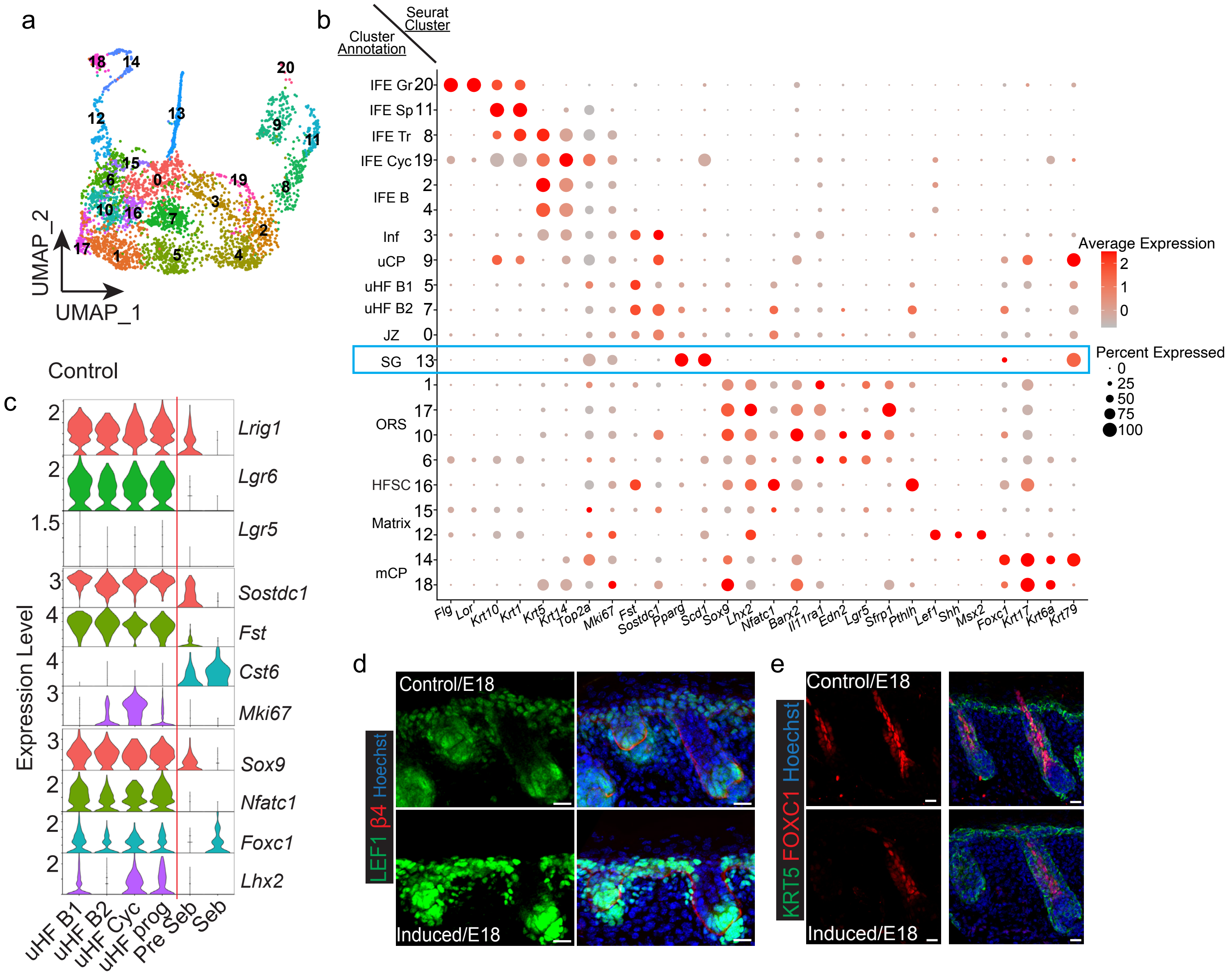

### Extended Data Figure 4

**Extended Data Figure 4**

a

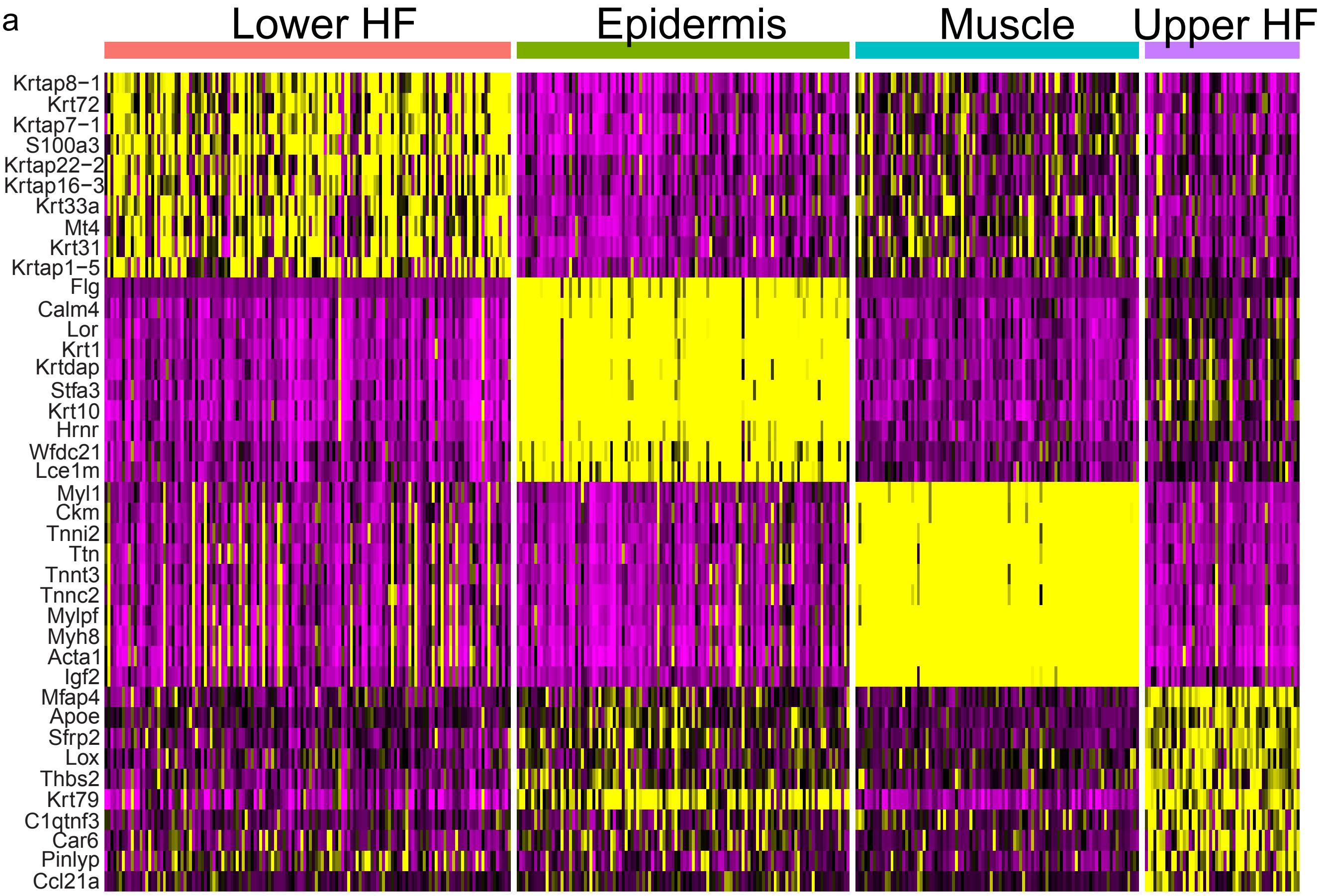


b

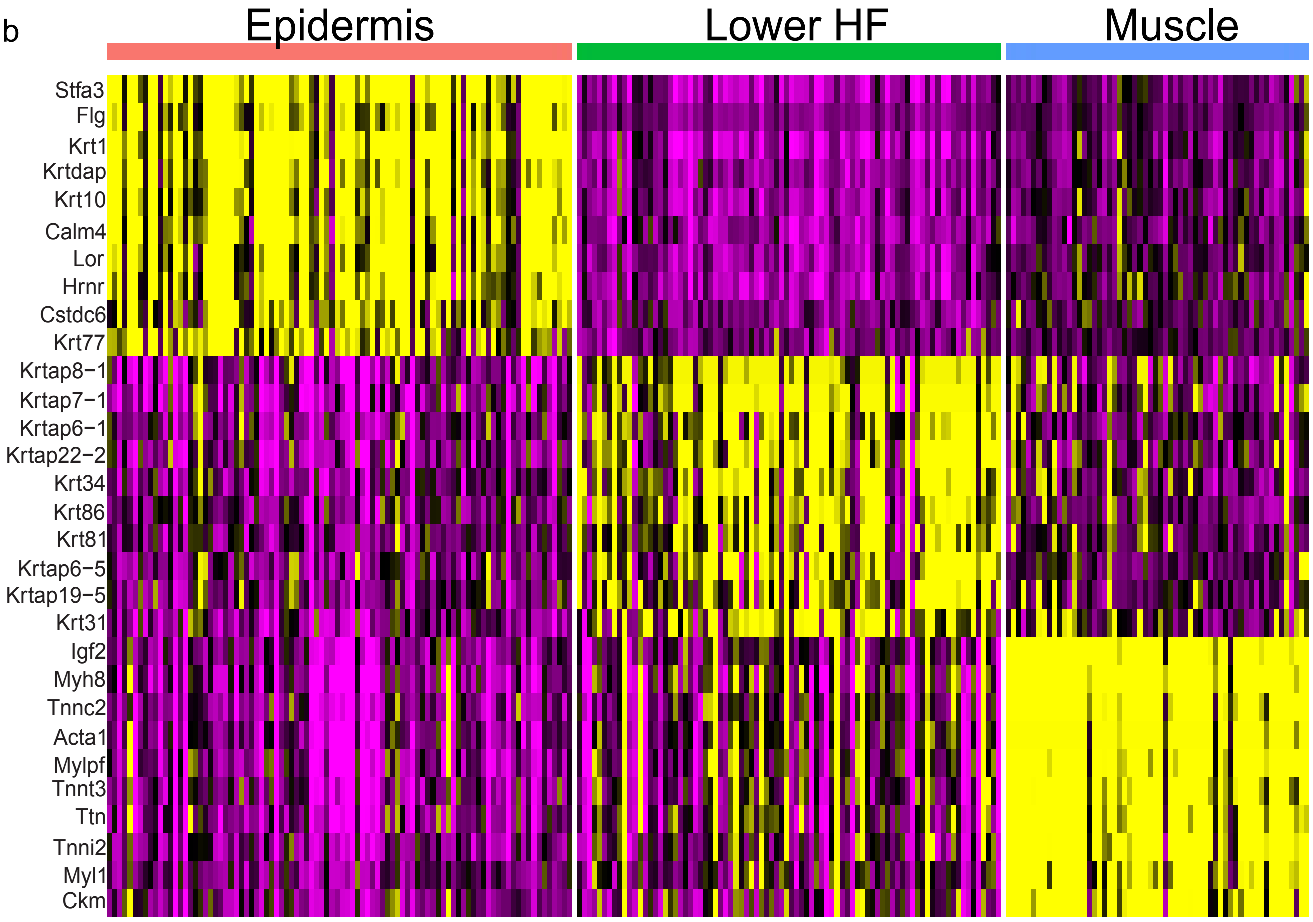

### Extended Data Figure 5

##### Extended Data Figure 5

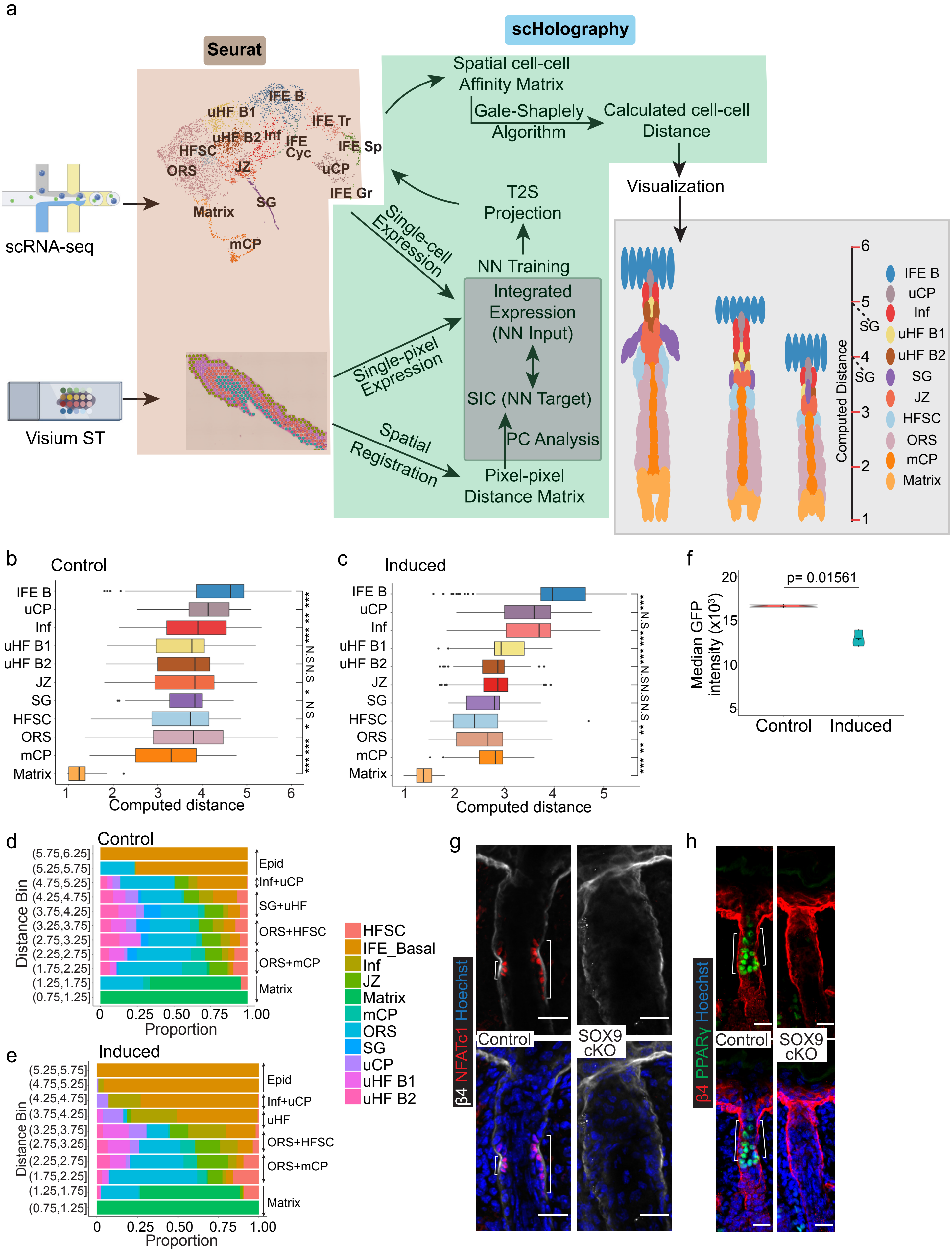

### Extended Data Figure 6

**Extended Data Figure 6**

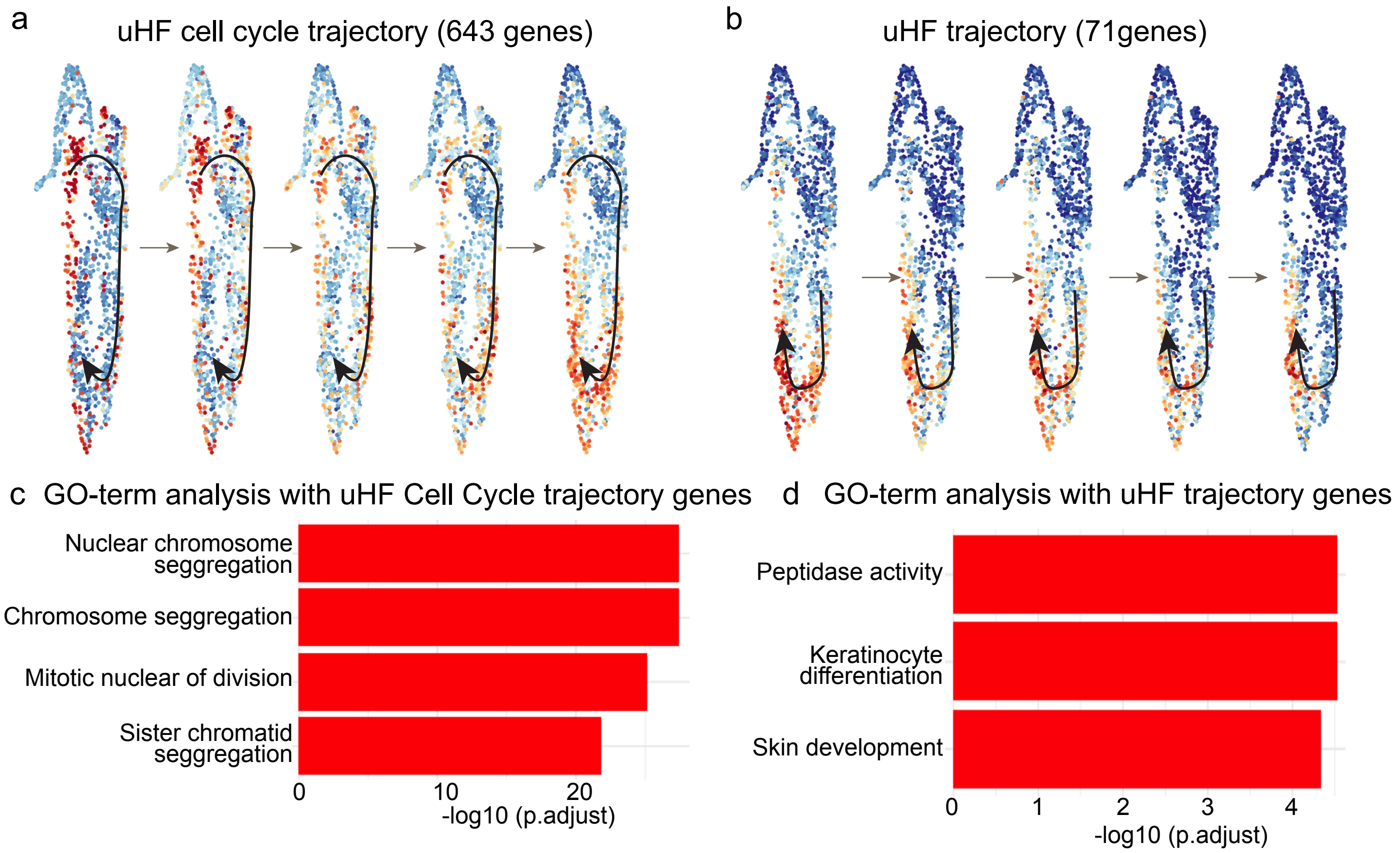

### Extended Data Figure 7

**Extended Data Figure 7**

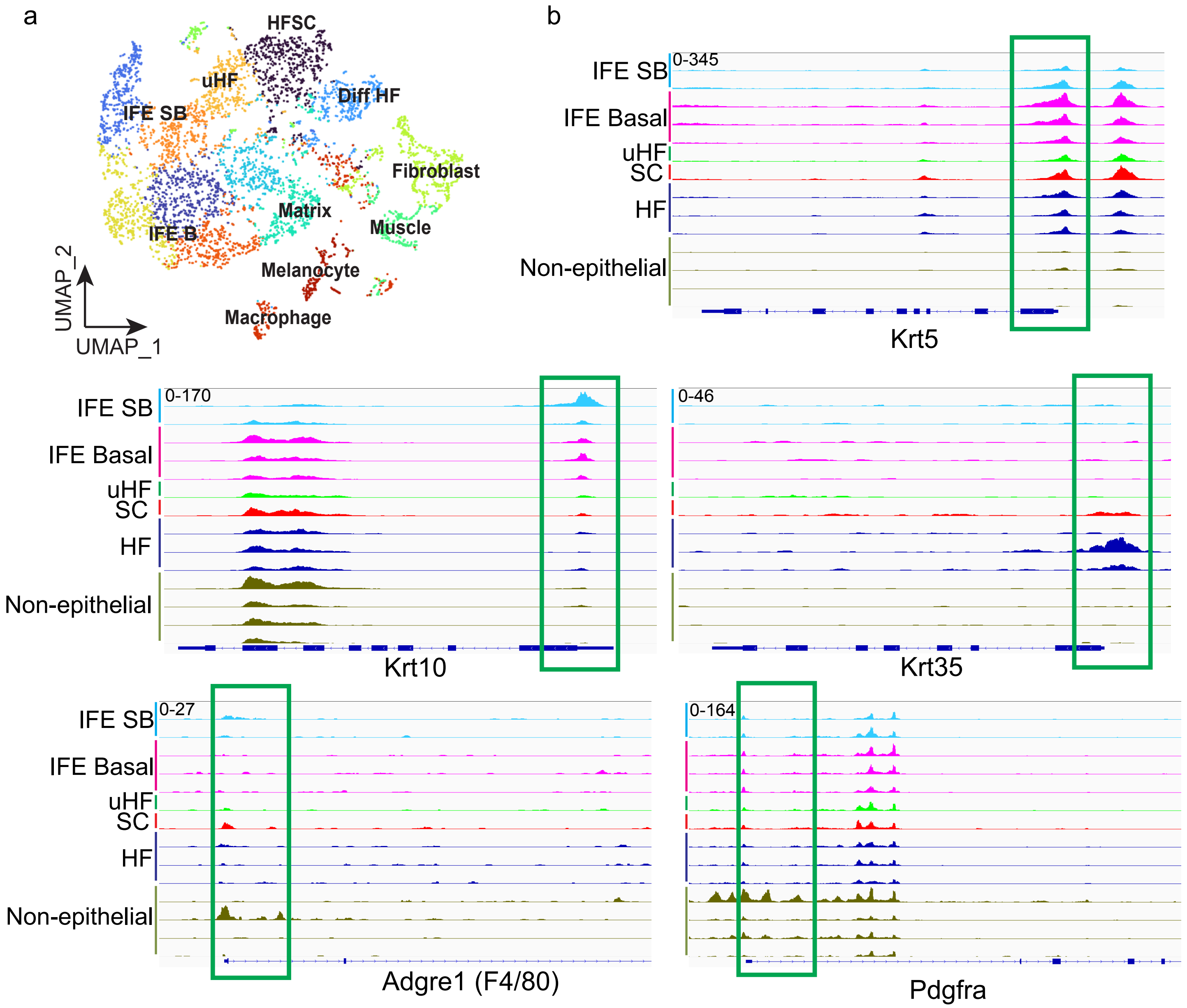
